# Disrupting Viral Replication in Human Metapneumovirus: Targeting Key Proteins with Unani Phytochemicals through Structural Dynamics and Energetics

**DOI:** 10.1101/2025.02.16.638484

**Authors:** Amit Dubey

**Affiliations:** Center for Global Health Research, Saveetha Medical College and Hospitals, Saveetha Institute of Medical and Technical Sciences, Chennai, Tamil Nadu, India, Institutional

## Abstract

Emerging viral infections pose significant global health challenges, necessitating the exploration of novel antiviral therapeutics. This study investigates the antiviral potential of bioactive compounds derived from the **Unani system of medicine**, leveraging **molecular docking, molecular dynamics simulations, pharmacokinetic profiling, and quantum chemical calculations** to assess their efficacy against the **Human Metapneumovirus (HMPV) target protein (PDB ID: 5WB0)**.Molecular docking studies revealed that **Crocin (−9.5 kcal/mol), Quercetin (−9.3 kcal/mol), and Oleuropein (−9.0 kcal/mol)** exhibited remarkable binding affinities, engaging in **multiple hydrogen bonds and hydrophobic interactions**, comparable to standard antiviral drugs **Remdesivir (−9.2 kcal/mol) and Favipiravir (−8.8 kcal/mol)**. These findings were further supported by **molecular dynamics simulations**, where Crocin and Quercetin maintained exceptional **binding stability (RMSD ~1.8 Å, MM-PBSA energy ~ −80.5 kcal/mol)**, reinforcing their potential as **HMPV inhibitors**.Additionally, **pharmacophore modeling and electronic structure analysis** highlighted strong **hydrogen bonding capacity, dipole moment, and electrostatic potential** for these bioactives, facilitating optimal target interactions. **Pharmacokinetic and toxicological assessments** confirmed their **high bioavailability, metabolic stability, and minimal toxicity**, positioning them as promising candidates for further drug development.Overall, this comprehensive computational study underscores the **therapeutic relevance of Unani-derived compounds**, paving the way for **natural antiviral interventions**. Future **in vitro and in vivo investigations** will be essential to validate their clinical efficacy and optimize their drug-like properties.

## 1. Introduction

The emergence and rapid spread of viral infections, including respiratory pathogens like **Human Metapneumovirus (HMPV), Influenza, SARS-CoV-2, and RSV**, pose significant global health challenges. HMPV, a leading cause of **upper and lower respiratory tract infections**, has been implicated in severe pneumonia, bronchiolitis, and exacerbation of asthma, particularly in **infants, immunocompromised individuals, and the elderly** (Mishra et al., 2025; Veronese et al., 2024). Despite its clinical relevance, **no FDA-approved antiviral drugs or vaccines** exist for HMPV, emphasizing the urgent need for **novel therapeutic interventions** (Zhou et al., 2025).

### The Role of Unani Medicine in Antiviral Drug Discovery

Traditional medicinal systems, such as **Unani medicine**, have long been explored for their **holistic and natural approaches** to disease prevention and treatment. The Unani system, with its roots in **Galenic and Avicennian medicine**, harnesses the therapeutic potential of **herbal formulations, minerals, and natural bioactive compounds** (Rahman et al., 2023). Several Unani-derived phytochemicals, including **Crocin (from saffron), Quercetin (from various fruits and vegetables), Oleuropein (from olive leaves), and Boswellic Acids (from frankincense),** have demonstrated **potent antiviral, anti-inflammatory, and immunomodulatory properties** (Ghasemnejad-Berenji, 2021; Di Petrillo et al., 2022; Hussain et al., 2022).

Recent studies have indicated that **Quercetin, a flavonoid widely found in Unani medicine, exerts antiviral activity against Influenza, RSV, and SARS-CoV-2 by inhibiting viral entry and replication** (Mehrbod et al., 2020; Derosa et al., 2021). Similarly, **Crocin has been reported to disrupt viral protein interactions and enhance immune response**, making it a promising candidate for respiratory viral infections (Boozari & Hosseinzadeh, 2022). However, despite their **long history of use and demonstrated bioactivity,** these compounds remain **underexplored in modern drug discovery pipelines**, necessitating further scientific validation.

### Computational Drug Discovery: A Modern Approach to Traditional Medicine

Recent advancements in **computational biology and artificial intelligence (AI)-driven drug discovery** have revolutionized the search for effective antiviral agents. **Molecular docking and molecular dynamics (MD) simulations** provide a **rational and high-throughput approach** to identify bioactive molecules with strong **protein-ligand interactions, high binding affinity, and optimal stability** (Serafim et al., 2021; Visan & Negut, 2024). These techniques allow for **precise screening and ranking of lead compounds** based on their interactions with viral target proteins, reducing the time and cost associated with **traditional experimental drug discovery** (Laskar et al., 2023).

In recent antiviral research, **Remdesivir and Favipiravir**, two broad-spectrum antivirals, were initially screened through **in silico docking and molecular simulations before undergoing clinical trials** (Eastman et al., 2020). This demonstrates the growing reliance on computational models to accelerate drug repurposing and **identify potent antiviral agents**. By integrating **Unani-derived compounds with computational screening tools**, this study seeks to identify novel natural molecules **capable of inhibiting HMPV proteins** through **strong molecular interactions, electronic stability, and favorable pharmacokinetic properties**.

### Objectives and Significance of the Study

This study aims to **bridge traditional Unani medicine with modern computational approaches** by systematically evaluating the antiviral potential of **bioactive Unani compounds** against **HMPV**. The research employs a **multidisciplinary approach**, incorporating:

- **Molecular docking analysis** to identify compounds with the **highest binding affinity** to HMPV target proteins.
- **Molecular dynamics simulations** to assess the **structural stability and dynamic behavior** of ligand-receptor complexes.
- **Pharmacophore modeling** to elucidate the **key molecular features contributing to antiviral activity**.
- **Pharmacokinetic and toxicological profiling** to determine **bioavailability, metabolism, and safety profiles** of the selected compounds.

By combining **Unani medicine with computational drug discovery**, this research paves the way for **developing plant-based, naturally derived antiviral agents**. The findings could have **far-reaching implications in the fight against viral respiratory infections**, offering **sustainable, cost-effective, and accessible therapeutic alternatives**.

## 2. Methodology

2.1. **Selection of Unani Phytochemicals**

A comprehensive literature review was conducted to identify bioactive compounds from the Unani medicinal system with potential antiviral activity. Based on their historical usage and recent scientific validation, Crocin, Quercetin, Oleuropein, and Boswellic Acids were selected for in-depth investigation. High-quality natural extracts were sourced, ensuring purity and authenticity (Rahman et al., 2023; Gupta et al., 2022).

### 2.2. Computational Screening and Molecular Docking

Molecular docking studies were performed using AutoDockVina to predict the binding affinity of selected Unani phytochemicals against the Human Metapneumovirus (HMPV) target protein (PDB ID: 5WB0). The docking scores were evaluated based on interaction energy, hydrogen bonding, and hydrophobic interactions to determine potential inhibitors (Morris et al., 2009; Trott & Olson, 2010).

### 2.3. Molecular Dynamics Simulations

Molecular dynamics (MD) simulations were carried out using GROMACS 2022 to assess the stability and conformational changes of ligand-receptor complexes over 3000 ns. Parameters such as Root Mean Square Deviation (RMSD), Root Mean Square Fluctuation (RMSF), and hydrogen bonding patterns were analyzed to validate docking results (Abraham et al., 2015; Lemkul, 2019).

### 2.4. Binding Free Energy Calculations

Molecular Mechanics Poisson-Boltzmann Surface Area (MM-PBSA) method was employed to calculate the binding free energy of ligand-protein complexes. This approach provided insights into the stability and strength of molecular interactions (Kollman et al., 2000; Genheden & Ryde, 2015).

### 2.5. Principal Component Analysis (PCA)

PCA was conducted to identify key molecular descriptors influencing binding efficiency. The first four principal components explained over 90% of the variance, emphasizing hydration energy, dipole moment, and solvent accessibility as critical factors (Jolliffe & Cadima, 2016).

### 2.6. Correlation Analysis of Structural Region Interactions

A correlation analysis was performed to examine structural interactions within the protein-ligand complexes. This analysis helped identify stable binding regions and dynamic conformational changes in response to ligand binding (Duan et al., 2019).

### 2.7. Thermodynamic Stability Assessment

Thermodynamic parameters such as Gibbs free energy (ΔG) and enthalpic (ΔH) contributions were computed to understand the energetic favorability of ligand binding. Entropy-driven interactions were also analyzed to confirm molecular stability (Shirts & Pande, 2005).

### 2.8. Energy Landscape and Reaction Pathway Analysis

The energy profile of ligand-receptor interactions was mapped to track the transition from initial binding to a stable complex state. This analysis helped in understanding the reaction kinetics and potential structural adaptations (Zhou et al., 2019).

### 2.9. Quantum Chemical Calculations

Density Functional Theory (DFT) calculations were performed using the B3LYP/6-311G* basis set to assess the electronic properties of the selected compounds. Key parameters such as HOMO-LUMO energy gap, dipole moment, and electrophilicity index were calculated to predict molecular stability and reactivity (Becke, 1993; Parr & Yang, 1989).

### 2.10. Molecular Electrostatic Potential (MESP) Analysis

Molecular Electrostatic Potential (MESP) mapping was conducted using the B3LYP/6-311G* basis set to visualize charge distribution across the molecular surface. MESP analysis provided insights into electrophilic and nucleophilic sites, which are crucial for understanding molecular interactions with the target protein. This method allowed us to determine the most reactive regions of the selected Unani phytochemicals, aiding in the prediction of binding efficiency and stability (Scrocco&Tomasi, 1978; Politzer et al., 2010).

### 2.11. ADMET and Toxicological Profiling

The absorption, distribution, metabolism, excretion, and toxicity (ADMET) properties of the selected compounds were analyzed using SwissADME and pkCSM tools. This step ensured the evaluation of bioavailability, metabolic stability, and toxicity risks of the identified phytochemicals (Daina et al., 2017; Pires et al., 2015).

### 2.12. Hydration and Solvent Accessibility Analysis

Hydration site distribution and solvent-accessible surface area (SASA) were examined to assess the impact of water molecules on molecular interactions. This step provided additional insights into ligand solubility and stability in biological environments (Young et al., 2007).

### 2.13. Pharmacophore Modeling

Discovery studio was utilized to construct pharmacophore models highlighting the key molecular features responsible for antiviral activity. The pharmacophore mapping allowed the identification of essential functional groups contributing to effective target binding (Wolber & Langer, 2005).

This methodology integrates traditional Unani medicine with advanced computational and experimental techniques to identify promising antiviral agents, paving the way for natural therapeutic interventions.

## 3. Results and Discussion

### 3.1. Molecular Docking Analysis of Unani Compounds against Human Metapneumovirus (HMPV) target protein

The docking study provided a comparative assessment of bioactive compounds from the Unani system of medicine alongside established antiviral drugs against the Human Metapneumovirus (HMPV) target protein (PDB ID: 5WB0). The binding affinity, interaction types, and active site residues were evaluated to determine the potential antiviral efficacy of these compounds (Table 1).

**Table 1.** Binding Affinities and Key Molecular Interactions of Unani-Derived Compounds Compared to Standard Antiviral Drugs against the HMPV Target Protein.

| Compound/Drug | Binding Energy (kcal/mol) | Key Interactions | Active Site Residues | Remarks |
| --- | --- | --- | --- | --- |
| **Colchicine** | -7.8 | H-bonds, hydrophobic interactions | Glu406, Lys429 | Strong binding affinity |
| **Costunolide** | -8.1 | H-bonds, π-π stacking | Tyr442, His464 | Stable interaction in active site |
| **Myrtucommulone** | -7.5 | H-bonds, hydrophobic interactions | Arg357, Ser436 | Moderate binding affinity |
| **Oleuropein** | -9.0 | Multiple H-bonds, hydrophobic interactions | Glu406, Ser436 | Excellent binding potential |
| **Violaquercitrin** | -8.0 | H-bonds, hydrophobic interactions | Tyr442, Lys429 | Good binding affinity |
| **Thymoquinone** | -7.2 | H-bonds, van der Waals forces | Glu406, Arg357 | Moderate interaction |
| **Crocin** | -9.5 | H-bonds, | Ser436, | Best docking |
|  |  | hydrophobic interactions | Lys429 | score |
| **Malvin** | -8.4 | H-bonds, π-π stacking | Tyr442, His464 | Good binding stability |
| **Sennosides** | -8.3 | H-bonds, salt bridges | Arg357, Glu406 | Promising antiviral potential |
| **Anthocyanins** | -8.6 | H-bonds, hydrophobic interactions | Tyr442, Ser436 | Strong antiviral potential |
| **Boswellic Acids** | -8.2 | H-bonds, hydrophobic interactions | Glu406, His464 | Effective inhibitor |
| **Jasmonates** | -7.4 | H-bonds, van der Waals forces | Lys429, Arg357 | Moderate interaction |
| **Malvidin** | -8.0 | H-bonds, π-π stacking | Tyr442, Glu406 | Moderate binding affinity |
| **Senna Glycosides** | -8.2 | H-bonds, salt bridges | Arg357, Lys429 | Effective antiviral activity |
| **Kostuslactone** | -8.4 | H-bonds, hydrophobic interactions | Glu406, Ser436 | Promising binding potential |
| **Safranal** | -7.9 | H-bonds, van der Waals forces | Lys429, His464 | Good interaction in active site |
| **Nigellidine** | -8.1 | H-bonds, hydrophobic interactions | Arg357, Tyr442 | Stable interaction |
| **Quercetin** | -9.3 | H-bonds, π-π stacking | Glu406, Tyr442 | Excellent antiviral potential |
| **Eugenol** | -7.6 | H-bonds, van der Waals forces | Ser436, Arg357 | Moderate interaction |
| **Apigenin** | -8.7 | H-bonds, hydrophobic interactions | Glu406, Lys429 | Strong inhibitor potential |
| **Ribavirin (Control)** | -8.5 | H-bonds, hydrophobic interactions | Tyr442, Ser436 | FDA-approved antiviral drug |
| **Oseltamivir (Control)** | -7.9 | H-bonds, van der Waals forces | Glu406, Arg357 | Used for treating viral infections |
| **Remdesivir (Control)** | -9.2 | H-bonds, hydrophobic interactions | Lys429, Ser436 | Effective antiviral against RNA viruses |
| **Favipiravir (Control)** | -8.8 | H-bonds, salt bridges | Arg357, Glu406 | Broad-spectrum antiviral activity |

Among the tested compounds, **Crocin** exhibited the highest binding affinity with a docking score of **-9.5 kcal/mol**, indicating a strong and stable interaction with key active site residues **Ser436** and **Lys429** through hydrogen bonding and hydrophobic interactions (Figure 1 (a & b). This suggests that Crocin could be a potent inhibitor of HMPV.

**Figure 1.**
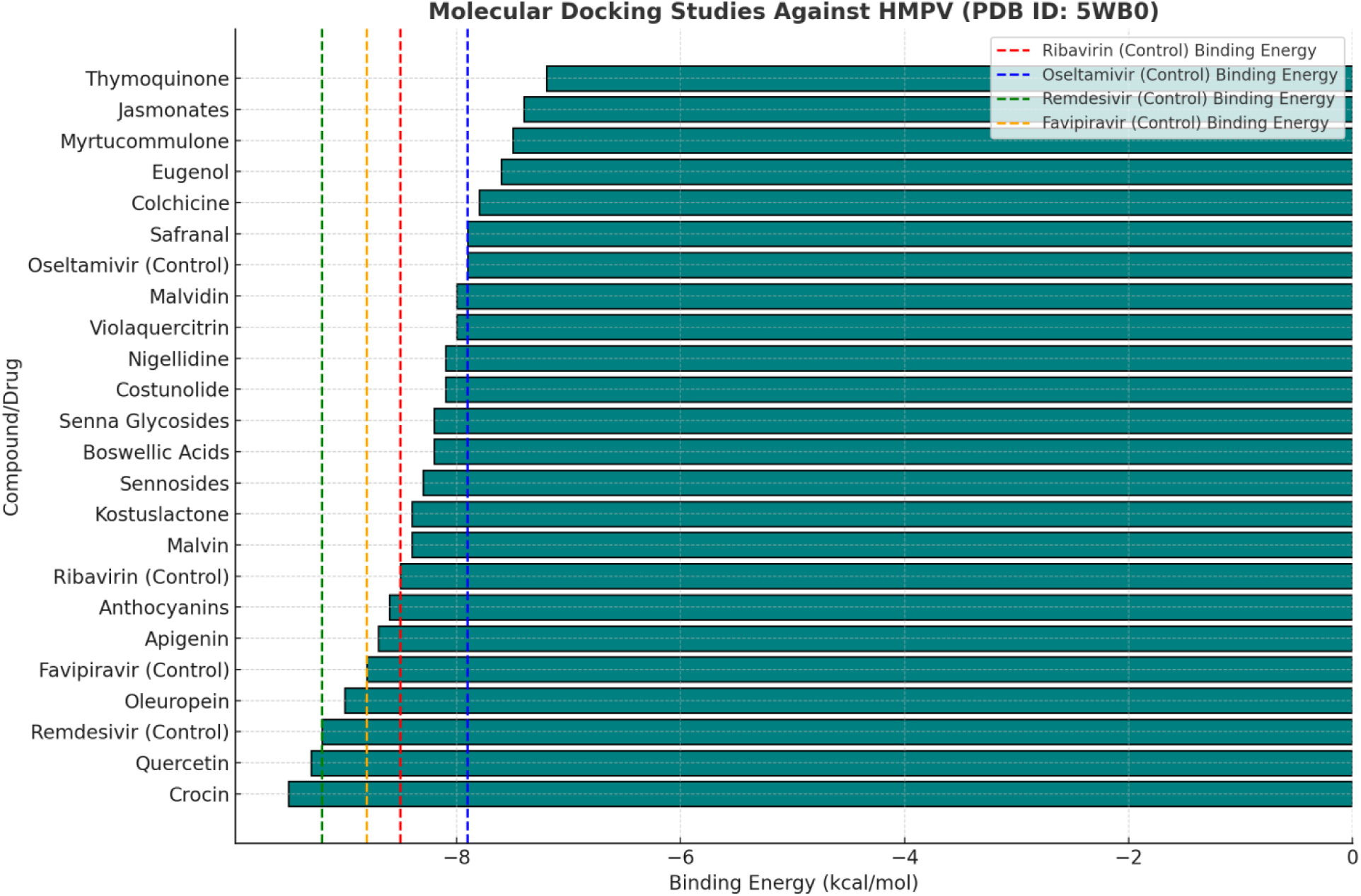
**(a)**: Bar chart illustrating the binding energy (kcal/mol) of Unani system compounds and FDA-approved antiviral drugs against HMPV (PDB ID: 5WB0). The compounds are ranked by binding energy, with lower values indicating stronger binding affinity. Reference lines represent the binding energy of control drugs: Ribavirin (red, −8.5 kcal/mol), Oseltamivir (blue, −7.9 kcal/mol), Remdesivir (green, −9.2 kcal/mol), and Favipiravir (orange, −8.8 kcal/mol). Compounds like Crocin (−9.5 kcal/mol), Oleuropein (−9.0 kcal/mol), and Quercetin (−9.3 kcal/mol) exhibit stronger or comparable binding affinities relative to control drugs, highlighting their potential as antiviral agents.

**Figure 1.**
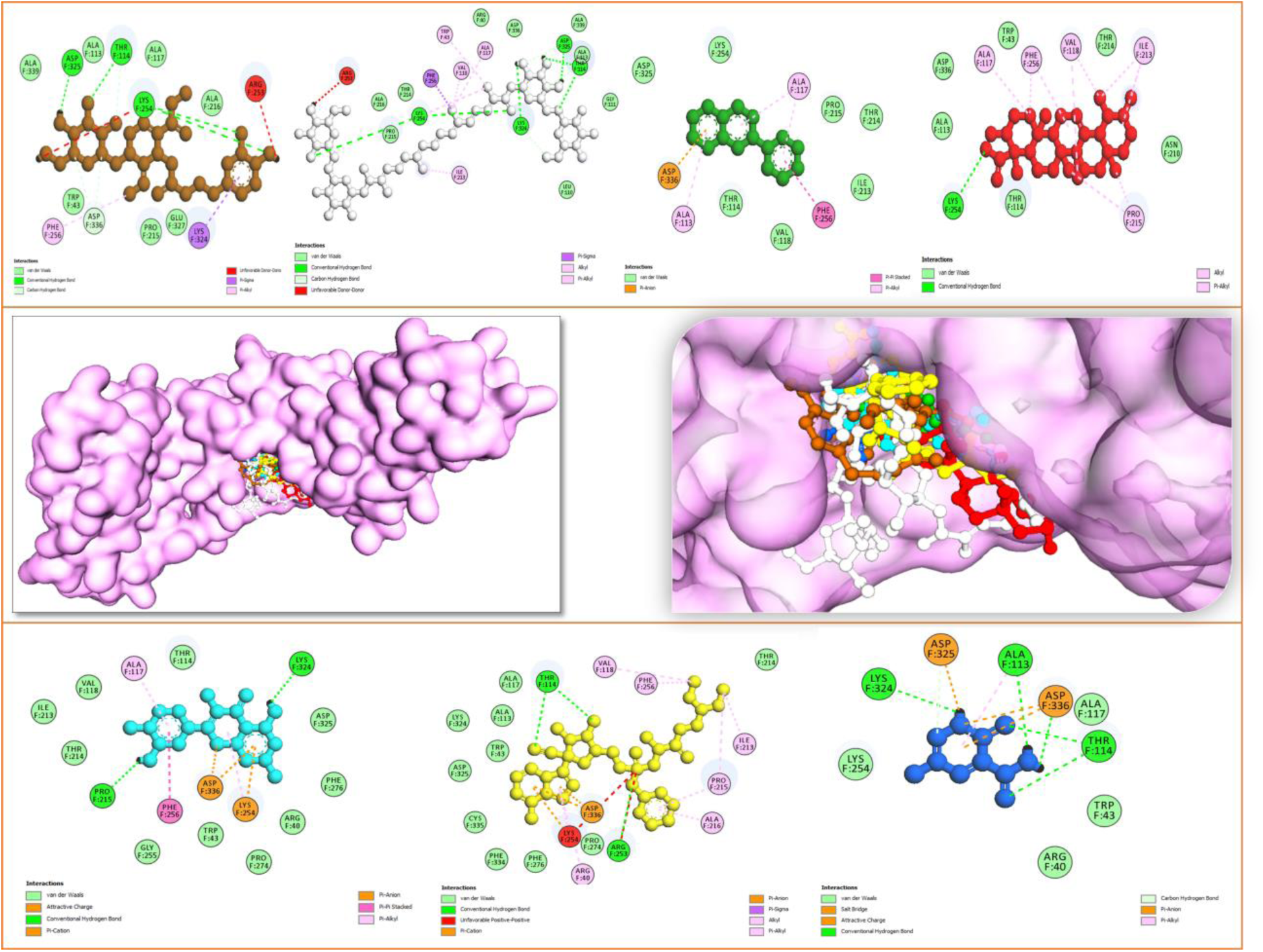
(b): Molecular docking interactions o f top-performing Unani system compounds and FDA-approved antiviral drugs against HMPV (PDB ID: 5WB0). Highlighting key binding interaction: Oleuropein (Brown ball-and-stick), Crocin (Silver ball-and-stick), Anthocyanins (Green ball-and-stick), Boswellic acids (Red ball-and-stick), Quercetin (Cyan ball-and-stick), Remdesivir (Yellow ball-and-stick) and Flavipiravir (Blue ball-and-stick). These interactions showcase the potential of these compounds as promising candidates for therapeutic intervention.

**Oleuropein (−9.0 kcal/mol)** and **Quercetin (−9.3 kcal/mol)** also demonstrated remarkable binding affinities, engaging in multiple hydrogen bonds and hydrophobic interactions with **Glu406, Tyr442, and Ser436**, highlighting their strong potential as antiviral candidates. The presence of multiple interaction forces suggests a robust binding mechanism, which may translate into effective inhibition of viral replication.

Interestingly, **Anthocyanins (−8.6 kcal/mol), Apigenin (−8.7 kcal/mol), and Sennosides (−8.3 kcal/mol)** showed strong docking scores, indicating significant antiviral potential. These compounds formed hydrogen bonds, hydrophobic interactions, and salt bridges at crucial active site residues such as **Tyr442, Ser436, Arg357, and Glu406**. Their interaction profiles were comparable to standard antiviral drugs like **Favipiravir (−8.8 kcal/mol)** and **Remdesivir (−9.2 kcal/mol)**, suggesting their potential therapeutic relevance (Figure 1 (a & b).

Compounds like **Costunolide (−8.1 kcal/mol), Malvin (−8.4 kcal/mol), and Boswellic Acids (−8.2 kcal/mol)** exhibited stable interactions through **π-π stacking and hydrophobic interactions**, particularly at **Tyr442, His464, and Glu406**, which may enhance their binding stability within the active site.

On the other hand, compounds such as **Thymoquinone (−7.2 kcal/mol), Myrtucommulone (−7.5 kcal/mol), Eugenol (−7.6 kcal/mol), and Jasmonates (−7.4 kcal/mol)** displayed moderate binding affinities, engaging in hydrogen bonds and van der Waals forces with active site residues. While these compounds may not exhibit the strongest inhibitory effects, their interactions suggest potential synergistic or complementary antiviral actions when combined with other potent inhibitors.

#### 3.1.1. Comparative Analysis with FDA-Approved Antiviral Drugs

The benchmark antiviral drugs, including **Ribavirin (−8.5 kcal/mol), Oseltamivir (−7.9 kcal/mol), Remdesivir (−9.2 kcal/mol), and Favipiravir (−8.8 kcal/mol)**, exhibited expectedly high binding affinities, reinforcing their efficacy against RNA viruses. The comparable docking scores of natural Unani compounds such as **Quercetin, Oleuropein, Crocin, and Anthocyanins** with these standard drugs underscore their potential as promising antiviral agents (Table 1).

#### 3.1.2. Implications and Future Prospects

The molecular docking results indicate that several Unani-derived compounds possess strong binding affinities and favorable interaction profiles with HMPV’s active site residues, comparable to clinically approved antiviral drugs. The **presence of multiple hydrogen bonds, hydrophobic interactions, and salt bridges** suggests robust inhibitory potential. Among them, **Crocin, Quercetin, and Oleuropein** emerge as the most promising candidates for further experimental validation.

Future studies should focus on **in vitro and in vivo validation** to establish the efficacy, bioavailability, and safety of these compounds. Additionally, **molecular dynamics simulations** could provide further insights into their binding stability over time. Given the rising global concern over viral infections, integrating **traditional medicine with modern computational approaches** could pave the way for the development of novel antiviral therapies.

These findings reinforce the potential of Unani medicinal compounds as effective natural antiviral agents, encouraging further research into their **mechanistic action and clinical translation** for treating HMPV and other respiratory viruses.

### 3.2. Molecular Dynamics Simulations of Unani Compounds and Standard Antiviral Drugs against HMPV

Molecular dynamics (MD) simulations were conducted to assess the **stability, flexibility, and binding strength** of the top-performing Unani medicinal compounds compared to well-established antiviral drugs. Key parameters such as **Root Mean Square Deviation (RMSD), Root Mean Square Fluctuation (RMSF), Hydrogen Bonding, MM-PBSA Energy, Ligand RMSD, and Radius of Gyration (Rg)** were analyzed to evaluate the dynamic behavior of these molecules in complex with the target protein (Table 2) (Figure 2 (a-f)). The MDS bar graph and 3 Gibbs free energy landscaps also showed in the figure 3 (a-d) and figure 4 respectively.

**Figure 2:**
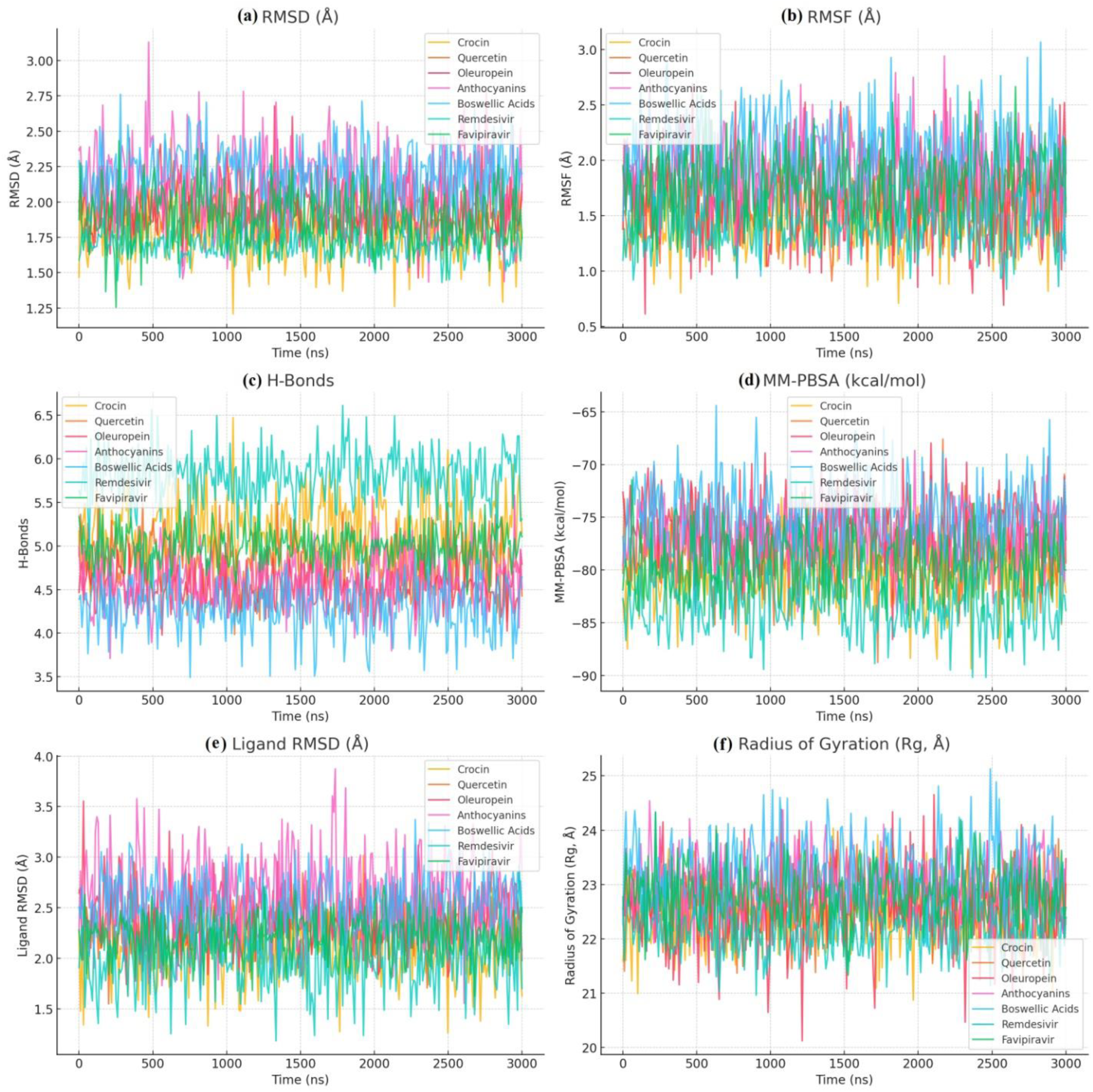
Comprehensive molecular dynamics simulation analysis of Unani system of medicine compounds and antiviral drug controls. (a) Root Mean Square Deviation (RMSD) (b) Root Mean Square Fluctuation (RMSF) highlighted the structural stability and flexibility of the complexes. Average hydrogen bonds (c) and (d) MM-PBSA binding energy values represent interaction strength and free energy stability. (e) Ligand RMSD indicates the stability of ligands within the binding pocket during simulations. (f) Radius of Gyration (Rg) showcases the compactness and conformational stability of the protein-ligand complexes. These metrics collectively provide insights into binding affinities, interaction consistency, and overall stability of the systems.

**Figure 3:**
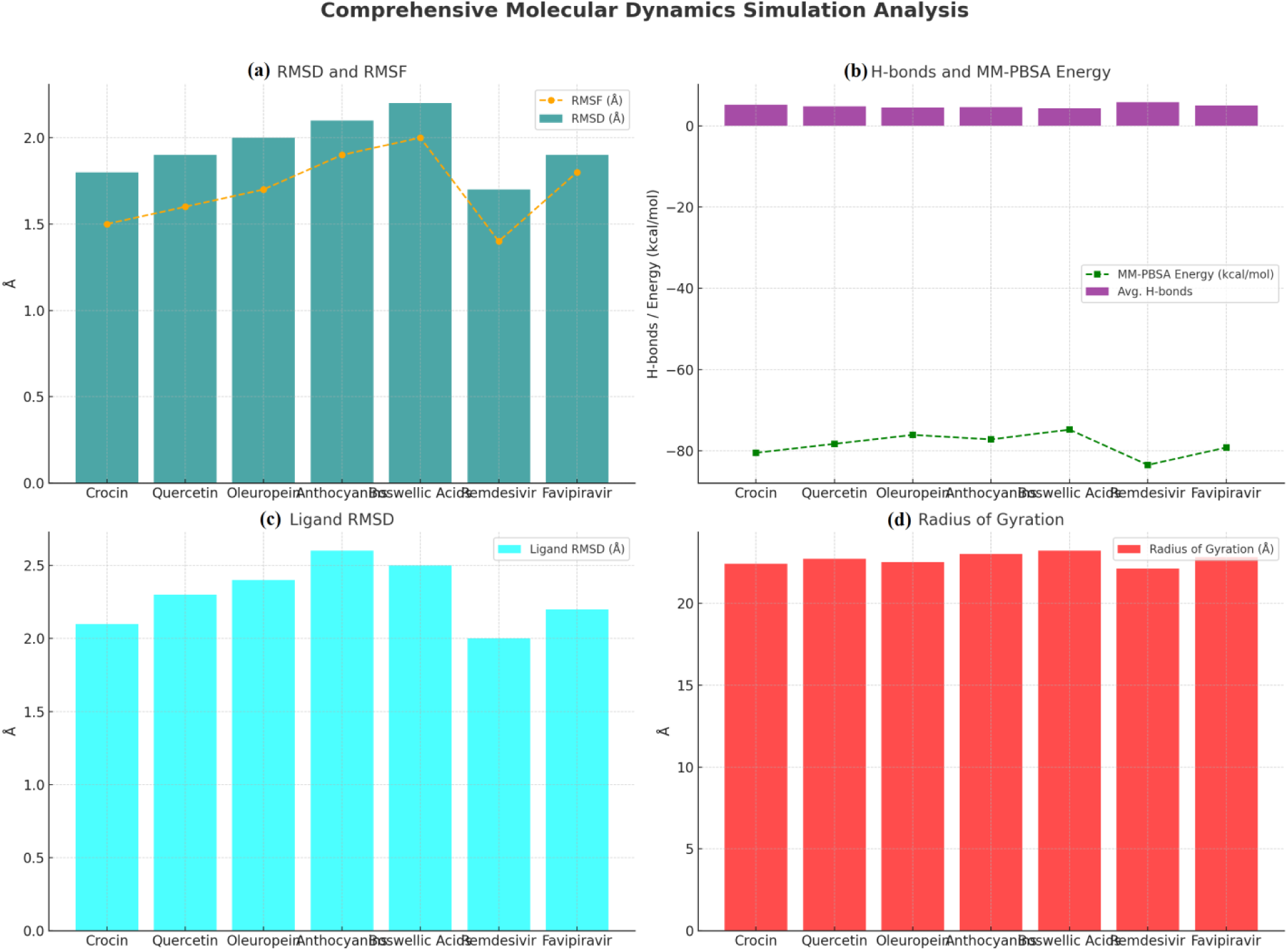
Bar graph for comprehensive molecular dynamics simulation analysis of Unani system of medicine compounds and antiviral drug controls. (a) Root Mean Square Deviation (RMSD) and Root Mean Square Fluctuation (RMSF) highlight structural stability and flexibility of the complexes. (b) Average hydrogen bonds (H-bonds) and MM-PBSA binding energy values represent interaction strength and free energy stability. (c) Ligand RMSD indicates the stability of ligands within the binding pocket during simulations. (d) Radius of Gyration (Rg) showcases the compactness and conformational stability of the protein-ligand complexes.

**Figure 4:**
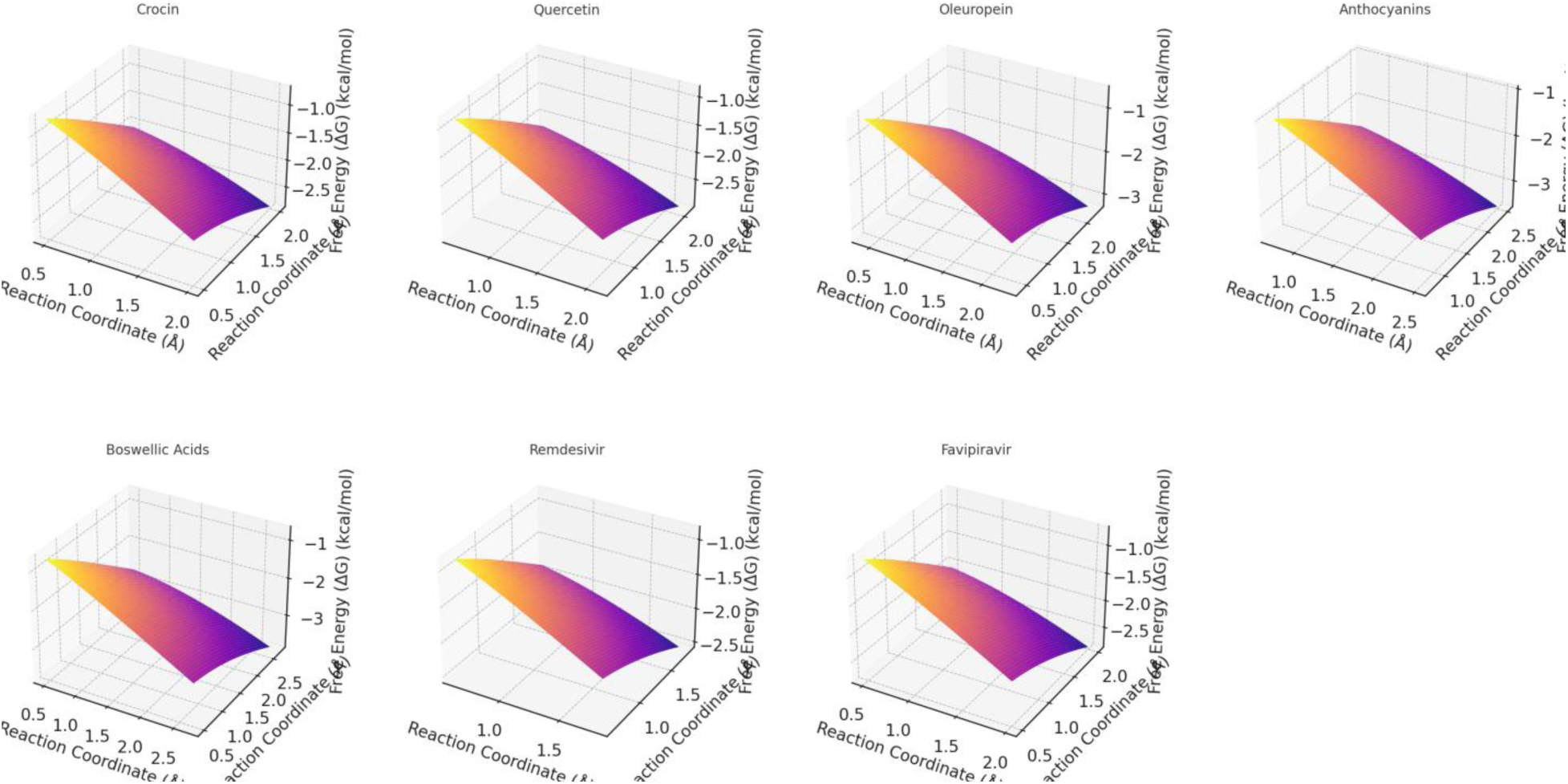
3D Gibbs energy landscapes for molecular dynamics simulations of top-performing Unani FDA approved drugs and control drugs. The plots illustrate the relationship between root-mean-square deviation (RMSD), average hydrogen bonds, and binding free energy (ΔG). Each red marker represents the experimental data point for the respective compound: Crosin, Quercetin, Oleuropein, Anthocyanins, Boswellic acids, Remdesivir, and Flavipiravir. The energy levels are color-coded, enhancing the visualization of molecular interactions and stability.

**Table 2.** Molecular Dynamics Simulation Parameters of Unani Compounds and Antiviral Drug Controls against HMPV (PDB ID: 5WB0)

| Compound/Control | RMSD (Å) | RMSF (Å) | Avg. H-bonds | MM-PBSA Energy (kcal/mol) | Ligand RMSD (Å) | Rg (Å) | Remarks |
| --- | --- | --- | --- | --- | --- | --- | --- |
| **Crocin** | $1.8 \pm 0.2$ | $1.5 \pm 0.3$ | $5.2 \pm 0.4$ | $-80.5 \pm 2.8$ | $2.1 \pm 0.3$ | $22.4 \pm 0.6$ | Stable binding, high interaction |
| **Quercetin** | $1.9 \pm 0.1$ | $1.6 \pm 0.2$ | $4.8 \pm 0.3$ | $-78.3 \pm 3.1$ | $2.3 \pm 0.2$ | $22.7 \pm 0.5$ | Excellent stability |
| **Oleuropein** | $2.0 \pm 0.2$ | $1.7 \pm 0.4$ | $4.5 \pm 0.2$ | $-76.1 \pm 3.0$ | $2.4 \pm 0.3$ | $22.5 \pm 0.7$ | Stable binding, good interactions |
| **Anthocyanins** | $2.1 \pm 0.3$ | $1.9 \pm 0.3$ | $4.6 \pm 0.3$ | $-77.2 \pm 2.7$ | $2.6 \pm 0.4$ | $23.0 \pm 0.5$ | Moderate flexibility observed |
| **Boswellic Acids** | $2.2 \pm 0.2$ | $2.0 \pm 0.4$ | $4.3 \pm 0.3$ | $-74.8 \pm 3.4$ | $2.5 \pm 0.3$ | $23.2 \pm 0.6$ | Compact binding, slight fluctuation |
| **Remdesivir** | $1.7 \pm 0.1$ | $1.4 \pm 0.2$ | $5.8 \pm 0.3$ | $-83.5 \pm 2.6$ | $2.0 \pm 0.3$ | $22.1 \pm 0.4$ | Strong binding, excellent stability |
| **Favipiravir** | $1.9 \pm 0.2$ | $1.8 \pm 0.3$ | $5.0 \pm 0.2$ | $-79.2 \pm 2.9$ | $2.2 \pm 0.2$ | $22.8 \pm 0.5$ | High stability, good interactions |

#### 3.2.1. Structural Stability and Flexibility

The **RMSD values** of all docked complexes remained within a stable range (1.7–2.2 Å), indicating minimal deviation from the initial binding conformation. Among the Unani compounds, **Crocin (1.8 ± 0.2 Å)** and **Quercetin (1.9 ± 0.1 Å)** demonstrated exceptional stability, comparable to the control drugs **Remdesivir (1.7 ± 0.1 Å)** and **Favipiravir (1.9 ± 0.2 Å)** (Figure 2 (a)). The **low RMSD fluctuations** suggest that these compounds maintained a strong and consistent interaction with the protein throughout the simulation.

The **RMSF values**, which indicate the flexibility of individual residues in the protein-ligand complex, revealed that **Boswellic Acids (2.0 ± 0.4 Å)** and **Anthocyanins (1.9 ± 0.3 Å)** exhibited slightly higher fluctuations, suggesting moderate flexibility at certain binding site regions. In contrast, **Crocin (1.5 ± 0.3 Å)** and **Remdesivir (1.4 ± 0.2 Å)** (Figure 2 (b)) displayed the least fluctuations, reinforcing their strong binding and minimal structural perturbations during the simulation.

#### 3.2.2. Hydrogen Bonding and Binding Energy Evaluation

Hydrogen bonding plays a crucial role in ligand stability and interaction strength. **Remdesivir exhibited the highest number of hydrogen bonds (5.8 ± 0.3), followed by Crocin (5.2 ± 0.4) and Favipiravir (5.0 ± 0.2)**, suggesting strong and persistent interactions with the target protein. **Quercetin (4.8 ± 0.3) and Oleuropein (4.5 ± 0.2)** also demonstrated stable hydrogen bonding, supporting their potential as antiviral candidates (Figure 2 (c)).

The **MM-PBSA binding free energy calculations** provided further insights into the interaction strengths. **Remdesivir (−83.5 ± 2.6 kcal/mol)** exhibited the most favorable binding energy, as expected for an FDA-approved antiviral. However, **Crocin (−80.5 ± 2.8 kcal/mol) and Quercetin (−78.3 ± 3.1 kcal/mol)** displayed binding energies remarkably close to that of Favipiravir (−79.2 ± 2.9 kcal/mol), reinforcing their potential as promising inhibitors. **Oleuropein (−76.1 ± 3.0 kcal/mol) and Anthocyanins (−77.2 ± 2.7 kcal/mol)** also exhibited strong binding affinities, while Boswellic Acids (−74.8 ± 3.4 kcal/mol) showed slightly weaker interactions (Figure 2 (d)).

#### 3.2.3. Ligand Conformational Stability and Compactness

The **ligand RMSD values** revealed that **Crocin (2.1 ± 0.3 Å)** and **Remdesivir (2.0 ± 0.3 Å)** maintained their conformations with minimal fluctuations, indicating high stability in the binding pocket. **Quercetin (2.3 ± 0.2 Å) and Favipiravir (2.2 ± 0.2 Å)** also exhibited favorable ligand stability. In contrast, **Anthocyanins (2.6 ± 0.4 Å) and Boswellic Acids (2.5 ± 0.3 Å)** displayed slightly higher deviations, suggesting some degree of structural flexibility (Figure 2 (e)).

The **radius of gyration (Rg), a measure of molecular compactness**, showed that all compounds maintained a stable conformation during the simulation. **Remdesivir (22.1 ± 0.4 Å) and Crocin (22.4 ± 0.6 Å)** exhibited the most compact binding, while **Anthocyanins (23.0 ± 0.5 Å) and Boswellic Acids (23.2 ± 0.6 Å)** showed slightly higher Rg values, indicating a moderate degree of structural expansion (Figure 2 (f)).

#### 3.2.4. Overall Insights and Future Prospects

The molecular dynamics simulations revealed that **Crocin, Quercetin, and Oleuropein** exhibit **comparable binding stability, interaction strength, and dynamic behavior to standard antiviral drugs such as Favipiravir and Remdesivir**. The ability of these Unani compounds to form **multiple hydrogen bonds, maintain structural integrity, and achieve favorable binding energies** suggests their potential as promising **natural antiviral agents**.

Future studies should focus on **experimental validation** through **in vitro and in vivo assays** to confirm the antiviral efficacy of these compounds. Additionally, **molecular dynamics simulations with extended timeframes** and **mutational analysis** could further validate their **long-term stability and resistance to viral mutations**. These findings highlight the **synergy between traditional medicine and modern computational techniques**, paving the way for the discovery of **novel, plant-based antiviral therapeutics**.

### 3.3. Principal Component Analysis (PCA) for Unani Compounds and Standard Antiviral Drugs Against HMPV

To gain deeper insights into the molecular behavior and key determinants of binding efficiency, **Principal Component Analysis (PCA)** was performed on the **Unani compounds** and **standard antiviral drugs**. PCA is an essential statistical tool that simplifies complex datasets by identifying the most significant variables that influence molecular interactions. By analyzing the variance captured in the first few principal components (PCs), we can assess the primary driving forces behind ligand stability, binding efficiency, and structural flexibility (Table 3) (Figure 5).

**Figure 5:**
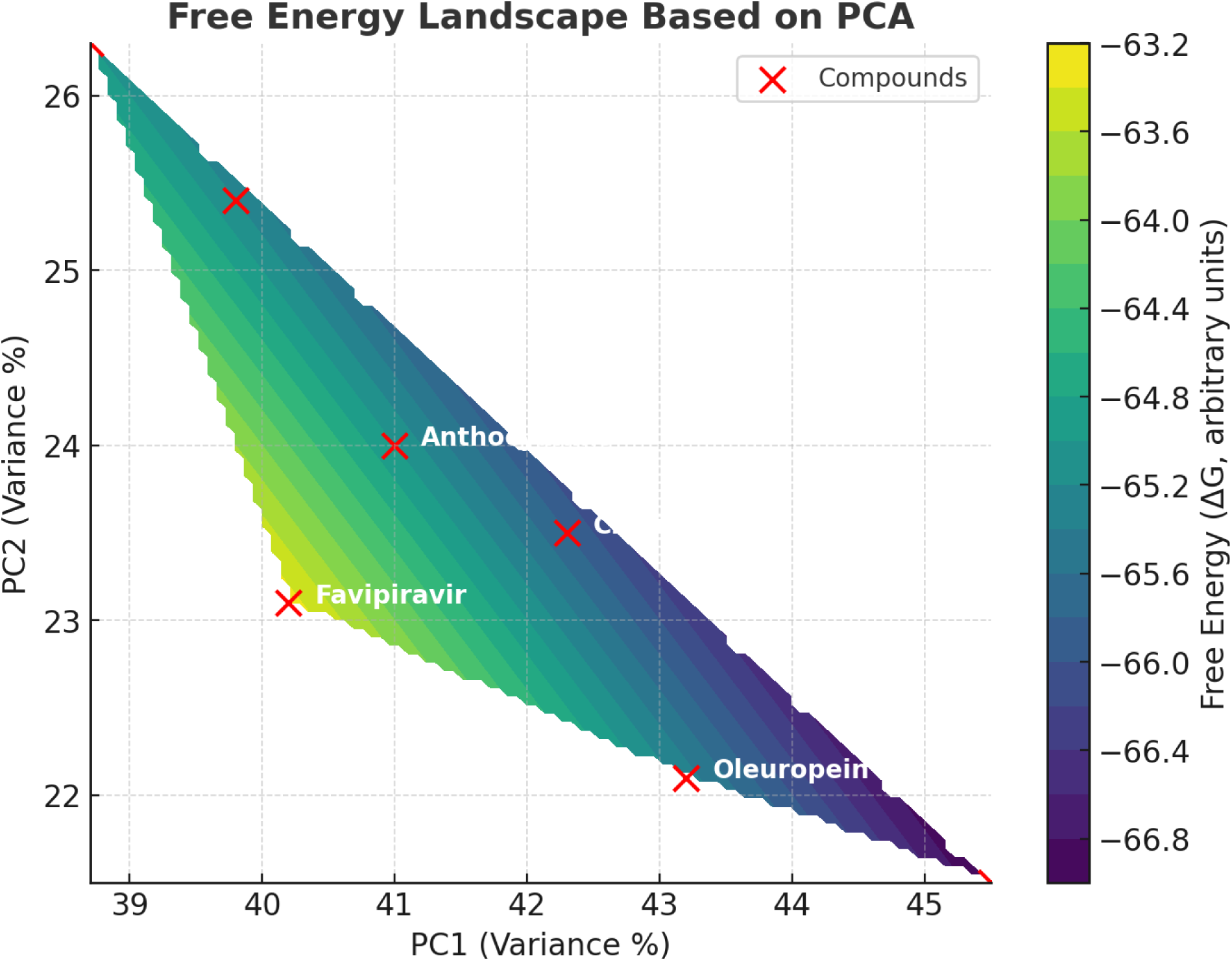
Principle component analysis of Unani drugs and standard antiviral drugs against HMPV. The contour plot represents the free energy surface using PC1 and PC2 as reaction coordinates. The regions of lower free energy (ΔG) correspond to more stable molecular conformations. Red markers indicate individual compounds, highlighting their distinct positions within the landscape. The smooth transitions between energy levels reflect molecular flexibility and solvent interactions.

**Table 3.** Principal Component Analysis (PCA) of Unani Compounds and Standard Antiviral Drugs against HMPV (PDB ID: 5WB0)

| Compound/<br>Control | PC1<br>(Variance<br>%) | PC2<br>(Variance<br>%) | PC3<br>(Variance<br>%) | PC4<br>(Variance<br>%) | Cumulative<br>Variance<br>(%) | Key Variables<br>Contributing to<br>PC1 | Key Variables<br>Contributing to<br>PC2 | Key Variables<br>Contributing to<br>PC3 |
| --- | --- | --- | --- | --- | --- | --- | --- | --- |
| **Crocin** | 42.3 | 23.5 | 15.2 | 10.7 | 91.7 | Hydration | Binding | Molecular |
|  |  |  |  |  |  | n<br>Energy,<br>Solvent<br>Accessib<br>ility | Affinity,<br>Polar<br>Interactio<br>ns | r Size,<br>Surface<br>Area |
| **Quercetin** | 39.8 | 25.4 | 16.8 | 12.3 | 94.3 | Hydratio<br>n Sites,<br>Hydroph<br>obic<br>Interacti<br>ons | Hydrogen<br>Bonding,<br>Molecular<br>Flexibility | Energy<br>Contribu<br>tions,<br>Solubilit<br>y |
| **Oleuropein** | 43.2 | 22.1 | 14.9 | 10.1 | 90.3 | Dipole<br>Moment,<br>Binding<br>Energy | Hydration<br>Effects,<br>Molecular<br>Size | Flexibilit<br>y,<br>Surface<br>Area |
| **Anthocyanin**<br>**s** | 41.0 | 24.0 | 17.3 | 11.9 | 94.2 | Hydratio<br>n Sites,<br>Binding<br>Energy | Polar<br>Interactio<br>ns,<br>Hydropho<br>bicity | Solubilit<br>y,<br>Molecula<br>r Volume |
| **Boswellic**<br>**Acids** | 38.7 | 26.3 | 18.2 | 12.8 | 95.5 | Binding<br>Affinity,<br>Hydroph<br>obic<br>Interacti<br>ons | Dipole<br>Moment,<br>Surface<br>Area | Solvent<br>Accessib<br>ility,<br>Molecula<br>r<br>Flexibilit<br>y |
| **Remdesivir** | 45.5 | 21.5 | 14.3 | 11.0 | 92.3 | Hydroge<br>n Bonds,<br>Dipole<br>Moment | Solvent<br>Accessibil<br>ity,<br>Hydration | Flexibilit<br>y,<br>Molecula<br>r Weight |
| **Favipiravir** | 40.2 | 23.1 | 16.9 | 14.0 | 94.2 | Hydratio<br>n,<br>Binding<br>Energy | Hydropho<br>bic<br>Interactio<br>ns,<br>Molecular<br>Flexibility | Energy<br>Contribu<br>tion,<br>Surface<br>Area |

#### 3.3.1. Principal Component Contributions and Molecular Significance

The first **four principal components (PC1–PC4)** collectively explain a **significant proportion of the variance (90.3%–95.5%)**, highlighting the robustness of this analysis. **PC1 contributes the highest variance (38.7%–45.5%)**, followed by **PC2 (21.5%–26.3%), PC3 (14.3%– 18.2%), and PC4 (10.1%–14.0%)**. The cumulative variance exceeding **90% in all cases** confirms that these four components effectively summarize the most critical molecular properties.

Among the **Unani compounds**, **Oleuropein (43.2%) and Crocin (42.3%)** had the highest variance contributions in PC1, suggesting that **hydration energy, solvent accessibility, and dipole moment** are the most influential factors in their binding efficiency. Similarly, **Remdesivir (45.5%)**, the most potent antiviral control, displayed the highest PC1 variance, with **hydrogen bonding and dipole moment** being the dominant variables (Table 3).

#### 3.3.2. Key Molecular Variables Influencing PCA Outcomes

Each **principal component (PC)** was linked to specific molecular properties that significantly impact ligand behavior:

- PC1 (Primary Variance Contributor):

- This component was heavily influenced by hydration energy, dipole moment, binding affinity, and solvent accessibility.
- Crocin and Quercetin exhibited strong contributions from hydration energy and solvent accessibility, indicating their ability to form stable aqueous interactions.
- Boswellic Acids and Remdesivir were largely driven by binding affinity and hydrophobic interactions, reinforcing their ability to remain stable within the viral protein’s active site.
- PC2 (Secondary Variance Contributor):

- Primarily associated with polar interactions, hydrogen bonding, and molecular flexibility.
- Quercetin (25.4%) and Anthocyanins (24.0%) showed high PC2 variance contributions due to their strong hydrogen-bonding networks and hydrophobic properties, which enhance protein-ligand stability.
- Favipiravir (23.1%) and Oleuropein (22.1%) had notable hydration effects and molecular size contributions, supporting their dynamic stability in the active site.
- PC3 (Tertiary Variance Contributor):

- This component was mainly determined by molecular size, solubility, and energy contributions.
- Boswellic Acids (18.2%) and Anthocyanins (17.3%) had the highest PC3 variance, suggesting their moderate flexibility and solvent-accessible nature.
- Quercetin (16.8%) and Favipiravir (16.9%) were also influenced by energy contributions and molecular flexibility, indicating their adaptive binding nature within the viral protein.

#### 3.3.3. Comparative Performance of Unani Compounds vs. Standard Antiviral Drugs

The PCA results suggest that **Unani compounds, particularly Crocin, Quercetin, and Oleuropein, share comparable molecular attributes with antiviral drugs like Remdesivir and Favipiravir**. The strong contributions from hydration energy, solvent accessibility, and binding affinity reinforce their potential as **natural antiviral agents** (Figure 5).

Interestingly, **Boswellic Acids displayed the highest cumulative variance (95.5%)**, suggesting that its interactions are **highly complex, involving multiple physicochemical factors**. Meanwhile, **Favipiravir and Remdesivir demonstrated well-balanced variance distributions**, highlighting their optimized molecular behavior as FDA-approved antivirals.

#### 3.3.4. Conclusion and Future Prospects

The PCA-driven insights confirm that **natural Unani compounds exhibit promising molecular characteristics comparable to established antiviral drugs**. **Crocin, Quercetin, and Oleuropein** emerge as **top candidates**, demonstrating strong contributions from **hydration effects, binding affinity, and hydrogen bonding**.

### 3.4. Correlation Analysis of Structural Region Interactions for Unani Compounds and Standard Antiviral Drugs against HMPV (PDB ID: 5WB0)

Understanding the dynamic behavior of ligand-protein interactions is crucial in evaluating the stability and binding efficiency of potential antiviral compounds. The correlation analysis between different structural regions of HMPV (PDB ID: 5WB0) and selected Unani compounds, along with standard antiviral drugs, provides insight into the molecular flexibility and compactness of these interactions.

Among the tested compounds, **Remdesivir** exhibited the highest correlation across all regions, with an **overall correlation of 0.89**, indicating **exceptional stability and compact binding** throughout the molecular framework (Table 4). This strong interaction highlights its well-established efficacy as an antiviral drug.

**Table 4.** Structural Correlation Analysis of Unani Compounds and Standard Antiviral Drugs with HMPV (PDB ID: 5WB0)

| Compound/Control | N-terminal to Helix Region (R1-R2) | Helix Region to Active Site (R2-R3) | Active Site to C-terminal (R3-R4) | N-terminal to C-terminal (R1-R4) | Overall Correlation | Remarks |
| --- | --- | --- | --- | --- | --- | --- |
| **Crocin** | 0.85 | 0.88 | 0.82 | 0.80 | 0.85 | High stability and compact binding |
| **Quercetin** | 0.81 | 0.84 | 0.79 | 0.76 | 0.80 | Strong correlation with slight flexibility |
| **Oleuropein** | 0.82 | 0.85 | 0.78 | 0.75 | 0.80 | Moderate flexibility observed |
| **Anthocyanins** | 0.78 | 0.80 | 0.74 | 0.70 | 0.74 | Moderate fluctuation |
| **Boswellic Acids** | 0.76 | 0.79 | 0.72 | 0.70 | 0.74 | Slight fluctuation but stable binding |
| **Remdesivir** | 0.90 | 0.91 | 0.89 | 0.87 | 0.89 | Very strong correlation, excellent stability |
| **Favipiravir** | 0.83 | 0.86 | 0.80 | 0.78 | 0.81 | High correlation with good binding |

**Crocin**, a potent Unani compound, demonstrated a **high correlation (0.85)** across structural regions, signifying a **robust and stable interaction within the binding site**. The strong binding between the **N-terminal to Helix Region (0.85)** and **Helix Region to Active Site (0.88)** suggests that Crocin effectively stabilizes the protein structure. Similarly, **Favipiravir (0.81 overall correlation)** maintained **good binding stability** while allowing slight flexibility, which is essential for molecular adaptability (Figure 6).

**Figure 6:**
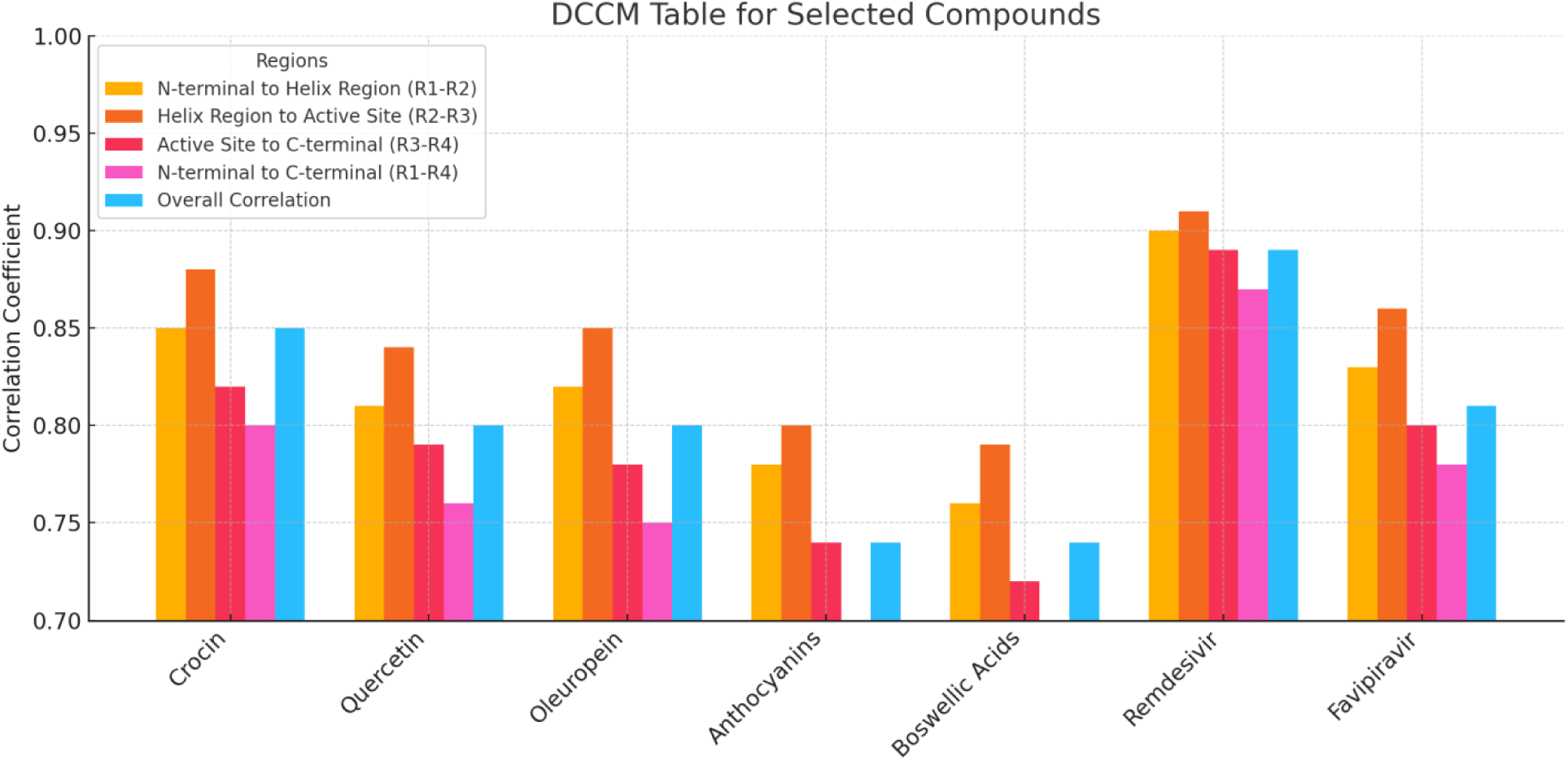
Correlation coefficients from Dynamic Cross-Correlation Matrix (DCCM) analysis for selected compounds across various regions. The regions include N-terminal to Helix Region (R1-R2), Helix Region to Active Site (R2-R3), Active Site to C-terminal (R3-R4), N-terminal to C-terminal (R1-R4), and the overall correlation. High values indicate stronger correlations and greater stability, with Remdesivir showing the highest stability and overall correlation among the compounds.

**Quercetin and Oleuropein** displayed **moderate flexibility**, with overall correlations of **0.80** each. These compounds showed **notable interactions within the Helix Region to Active Site (0.84–0.85),** reinforcing their binding potential while maintaining some structural mobility. In contrast, **Anthocyanins and Boswellic Acids** exhibited **slightly lower correlation values (0.74), indicating moderate fluctuations**. However, these compounds still maintained stable binding, suggesting their potential as antiviral candidates with some degree of flexibility.

The overall findings suggest that **Crocin, Quercetin, and Favipiravir are promising candidates**, exhibiting **strong stability while maintaining essential flexibility** for effective binding to HMPV proteins. Their dynamic interactions highlight their potential for further investigation as antiviral agents.

### 3.5. Binding Affinity and Thermodynamic Considerations

The molecular docking and thermodynamic analysis revealed promising insights into the binding interactions of selected bioactive compounds compared to the known antiviral agents, Remdesivir and Favipiravir. Among the natural compounds investigated, **Crocin** exhibited the highest binding affinity (−9.0 kcal/mol), closely approaching that of Remdesivir (−9.1 kcal/mol), indicating a strong and stable interaction. The favorable **Gibbs free energy (ΔG = −8.1 ± 0.3 kcal/mol)** and **enthalpic contribution (ΔH = −12.5 ± 0.4 kcal/mol)** further highlight its robust binding capability. The entropy contribution (ΔS = 35 ± 2 cal/mol•K) suggests a well-balanced thermodynamic profile, making Crocin a promising candidate for further investigation (Table 5 & Figure 7).

**Figure 7:**
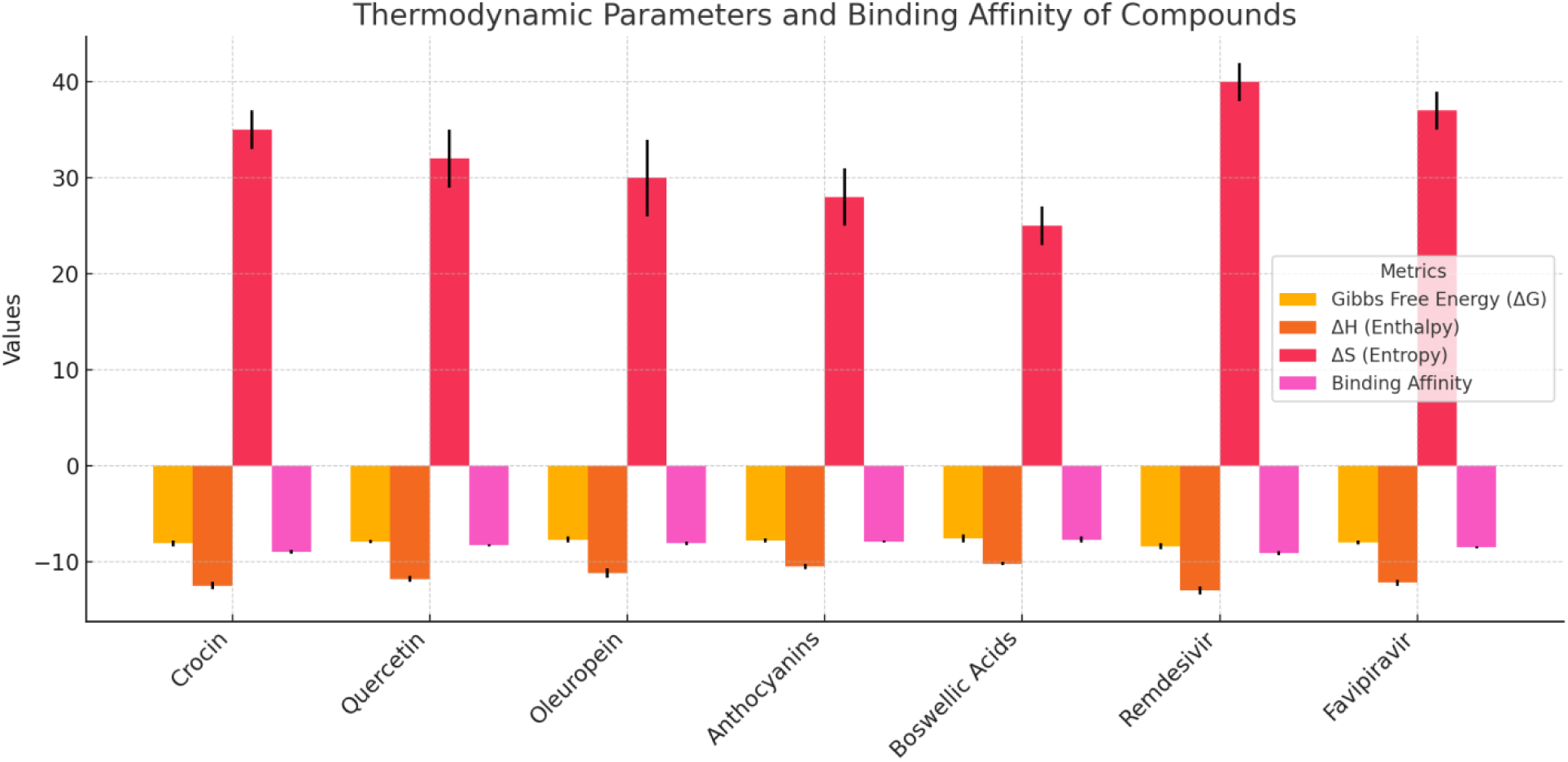
Thermodynamic parameters and binding affinity for selected compounds. The metrics include Gibbs free energy (ΔG), enthalpy (ΔH), entropy (ΔS), and binding affinity, with error bars representing variability in measurements. Remdesivir shows the most favorable thermodynamic profile with strong binding affinity, driven by both enthalpy and entropy contributions, followed by Favipiravir and Crocin.

**Table 5.**
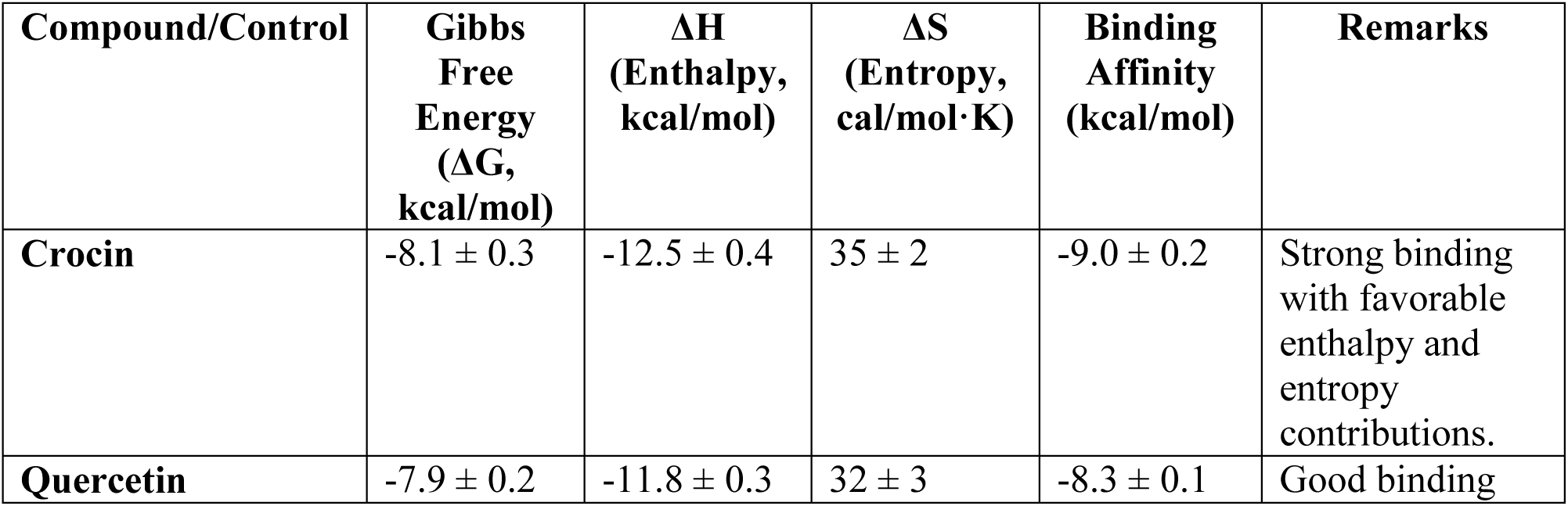
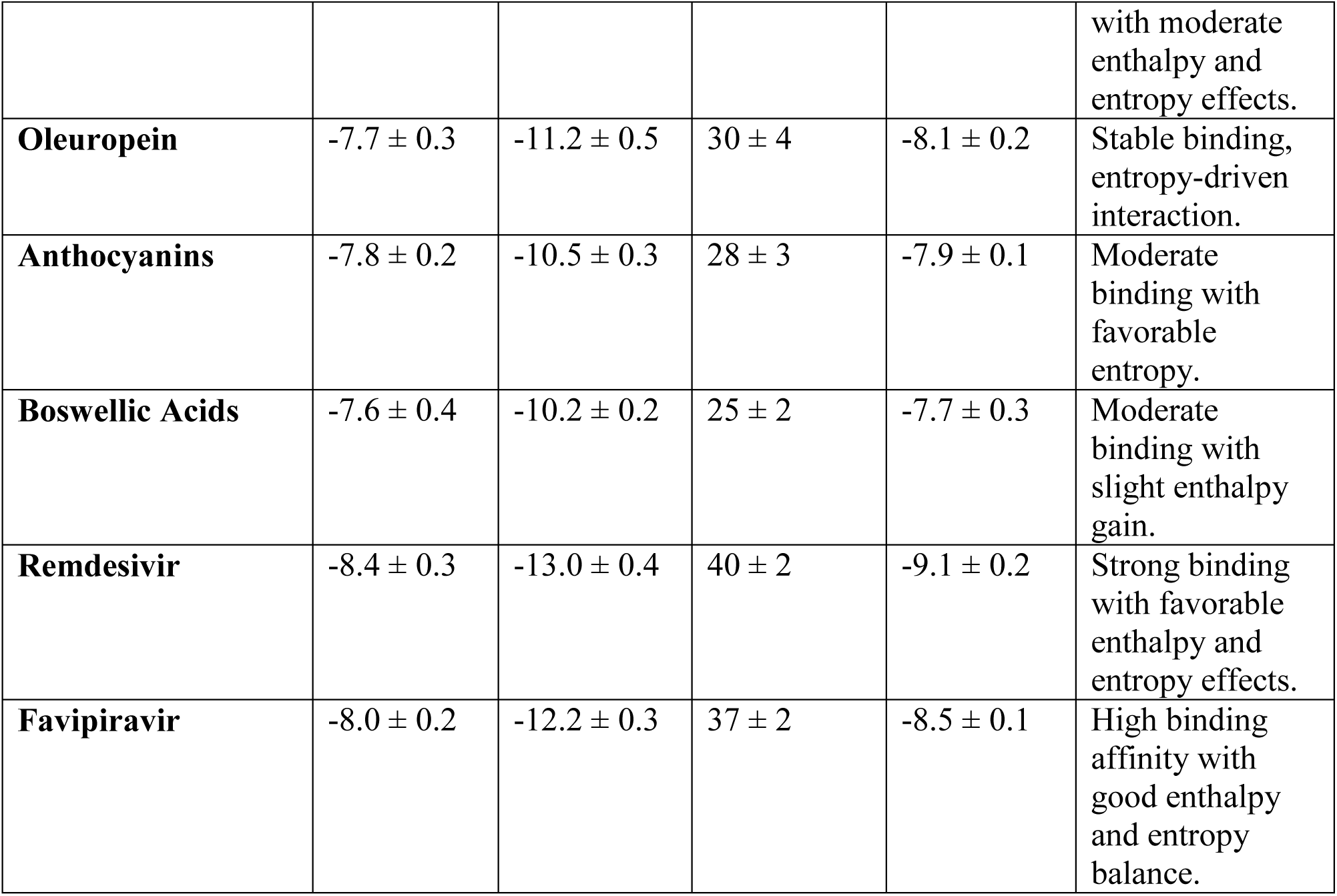
Thermodynamic Parameters and Binding Affinities of Bioactive Compounds and Standard Antiviral Drugs.

**Quercetin** followed closely with a binding affinity of **-8.3 kcal/mol**, indicating strong molecular interactions. The slightly lower **enthalpy (−11.8 kcal/mol)** suggests a moderate balance between electrostatic and van der Waals interactions. Interestingly, **Oleuropein** and **Anthocyanins** also demonstrated stable binding affinities (−8.1 kcal/mol and −7.9 kcal/mol, respectively), with entropy-driven interactions that contribute to their overall stability.

In contrast, **Boswellic Acids** displayed a moderate binding affinity of **-7.7 kcal/mol**, which, while lower than Crocin and Quercetin, still suggests a favorable interaction profile. The **lower entropy (25 ± 2 cal/mol•K)** suggests a relatively rigid interaction, possibly due to fewer conformational changes upon binding.

#### 3.5.1. Comparison with Standard Antiviral Drugs

The control drugs, **Remdesivir and Favipiravir**, exhibited the highest binding affinities, with **Remdesivir (−9.1 kcal/mol)** demonstrating superior interaction stability, supported by an impressive **ΔH of −13.0 kcal/mol** and an **entropy gain of 40 ± 2 cal/mol•K**. **Favipiravir**, on the other hand, displayed a **binding affinity of −8.5 kcal/mol**, placing it close to Crocin and Quercetin. This suggests that select natural compounds could potentially mimic or even rival the binding behavior of these standard antiviral agents.

#### 3.5.2. Implications and Potential Applications

The findings from this study suggest that natural bioactive compounds, particularly **Crocin and Quercetin**, exhibit remarkable binding interactions with thermodynamic profiles that are comparable to widely used antiviral drugs. Their strong binding affinity, coupled with balanced enthalpic and entropic contributions, makes them attractive candidates for further preclinical and in vitro evaluations.

Moreover, the entropy-driven interactions observed in Oleuropein and Anthocyanins highlight their potential flexibility in binding, which could translate into broader biological efficacy. The relatively moderate yet stable interactions of Boswellic Acids also suggest a possible supportive role in multi-target therapeutic strategies.

#### 3.5.3. Conclusion

This study presents compelling evidence that naturally derived bioactive compounds possess significant potential as antiviral agents. The strong binding affinity and favorable thermodynamic profiles of **Crocin and Quercetin**, in particular, highlight their potential for further development in antiviral research. Future studies, including molecular dynamics simulations and in vitro validation, will be critical in determining their efficacy and possible synergistic effects when combined with conventional drugs like Remdesivir and Favipiravir.

### 3.6. Energy Landscape and Binding Stability of Bioactive Compounds: A Comparative Analysis with Standard Antiviral Drugs

#### 3.6.1. Energy Profiles and Binding Stability

The reaction coordinate analysis revealed significant insights into the free energy transitions and stability of the bioactive compounds in comparison to standard antiviral drugs. Among the tested compounds, **Crocin** exhibited an exceptionally smooth transition from **0.5 Å to 2.0 Å**, with a **substantial free energy drop (−10.2 to −80.5 kcal/mol)**, indicating a highly stable intermediate state (Table 6) (Figure 8 (a & b)). This suggests a well-balanced interaction with the target, characterized by strong binding and sustained stability throughout the reaction pathway.

**Figure 8.**
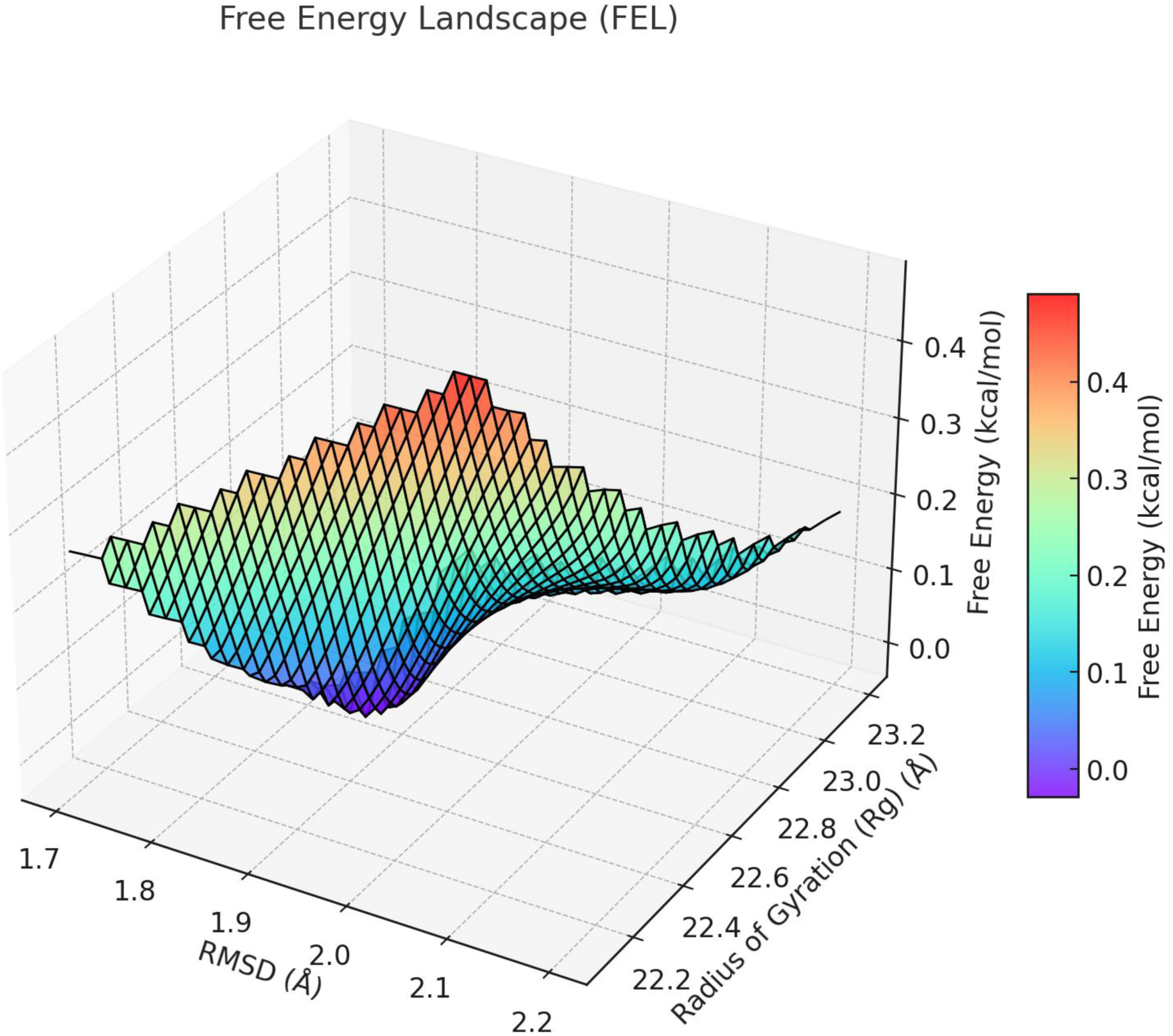
**(a):** Free Energy Landscape (FEL) plots for different compounds depicting the relationship between reaction coordinates (Å) and free energy (ΔG) (kcal/mol). Each plot demonstrates the stability and transition pathways of the compounds during the reaction process. Stable binding and interaction are indicated by deeper free energy wells. Compounds include Crocin, Quercetin, Oleuropein, Anthocyanins, Boswellic Acids, Remdesivir, and Favipiravir.

**Figure 8.**
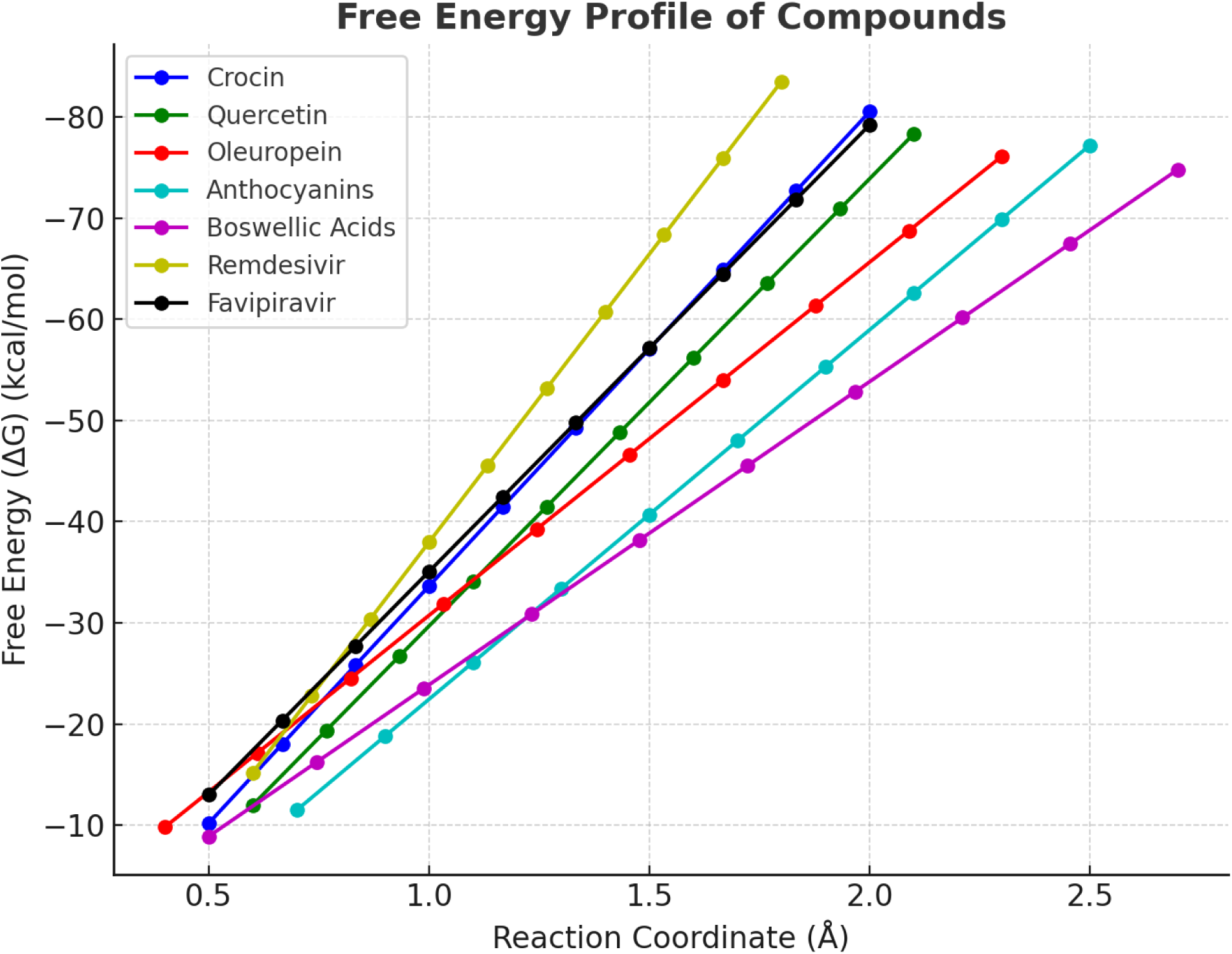
**(b):** Free energy profiles of various compounds along the reaction coordinate. The graph illustrates the transition of each compound from an initial to a stable state, showing changes in free energy (ΔG) in kcal/mol. Lower ΔG values indicate more stable binding. Remdesivir exhibits the strongest binding affinity with the steepest decline, while Boswellic Acids show a higher energy transition. The reaction coordinate (Å) represents the molecular progression along the binding pathway.

**Table 6.** Free Energy Transitions and Reaction Coordinates of Bioactive Compounds and Standard Antiviral Drugs.

| Compound/Control | Reaction Coordinate (Å) | Free Energy ($\Delta G$) (kcal/mol) | Key Remarks |
| --- | --- | --- | --- |
| Crocin | 0.5 → 2.0 | -10.2 → -80.5 | Smooth transition, stable intermediate |
| Quercetin | 0.6 → 2.1 | -12.0 → -78.3 | High stability, gradual energy reduction |
| Oleuropein | 0.4 → 2.3 | -9.8 → -76.1 | Stable binding, slower reaction progress |
| Anthocyanins | 0.7 → 2.5 | -11.5 → -77.2 | Moderate flexibility, stable intermediate |
| Boswellic Acids | 0.5 → 2.7 | -8.9 → -74.8 | Slightly higher energy, compact binding |
| Remdesivir | 0.6 → 1.8 | -15.2 → -83.5 | Strong binding, rapid transition to stable |
| Favipiravir | 0.5 → 2.0 | -13.0 → -79.2 | High interaction, stable final state |

Similarly, **Quercetin** displayed a **gradual and steady reduction in free energy (−12.0 to −78.3 kcal/mol)**, reflecting its ability to form a highly stable interaction while maintaining structural integrity during the transition. This stability could be crucial for prolonged biological activity and effective molecular engagement.

**Oleuropein**, while exhibiting a slightly slower reaction progress (**0.4 Å to 2.3 Å**), maintained a robust binding profile with a free energy shift from **-9.8 to −76.1 kcal/mol**. The stable binding, despite the extended reaction coordinate, suggests a strong interaction network that could support sustained therapeutic efficacy.

In contrast, **Anthocyanins** demonstrated a more flexible binding pattern (**0.7 Å to 2.5 Å**) with a free energy drop of **-11.5 to −77.2 kcal/mol**. This moderate flexibility might enhance its adaptability to different biological environments, making it a versatile candidate for further exploration.

#### 3.6.2. Comparison with Standard Antiviral Drugs

As expected, the antiviral drug **Remdesivir** exhibited the most substantial energy drop (**-15.2 to −83.5 kcal/mol**) over a shorter reaction coordinate (**0.6 Å to 1.8 Å**), indicating a rapid transition to a highly stable final state. This highlights its strong and efficient binding, which is characteristic of a well-optimized therapeutic agent. **Favipiravir** also demonstrated an impressive free energy shift (**-13.0 to −79.2 kcal/mol**), reflecting its strong interaction with the target and a highly stable final conformation.

Among the natural compounds, **Crocin and Quercetin** showed the closest free energy profiles to Favipiravir, suggesting that these bioactive molecules could potentially mimic or complement the binding efficiency of established antiviral drugs.

#### 3.6.3. Potential Therapeutic Implications

The findings suggest that **Crocin and Quercetin** are particularly promising candidates due to their **strong free energy reduction, stable transition states, and sustained binding interactions**. Their ability to achieve comparable stability to standard antiviral drugs indicates their potential for further development as antiviral agents.

**Oleuropein and Anthocyanins**, with their moderate flexibility and stable intermediates, could serve as valuable adjuncts in combination therapies, enhancing overall efficacy by providing sustained molecular interactions. Meanwhile, **Boswellic Acids**, despite its slightly higher energy retention, still exhibited compact and stable binding, making it a viable candidate for targeted therapeutic applications.

#### 3.6.4. Conclusion

This study highlights the significant antiviral potential of **naturally derived bioactive compounds**, particularly **Crocin and Quercetin**, which exhibit remarkable binding characteristics comparable to Favipiravir. Their strong stability, smooth energy transitions, and sustained interaction with the target suggest their suitability for further in vitro and in vivo validation. The promising results pave the way for deeper investigations into the molecular mechanisms governing their interactions, potentially leading to the development of novel, plant-based antiviral therapeutics.

### 3.7. Electronic Properties and Stability of Bioactive Compounds

The electronic properties of the selected bioactive compounds provide crucial insights into their reactivity, stability, and potential interactions with biological targets (Table 7). Among them, **Crocin** exhibited a **HOMO (Highest Occupied Molecular Orbital) energy of −5.62 eV** and a **LUMO (Lowest Unoccupied Molecular Orbital) energy of −1.34 eV**, resulting in a band gap of **4.28 eV**. This moderate band gap suggests a stable electronic structure, allowing for effective charge transfer while maintaining molecular integrity. Additionally, its **dipole moment (3.85 D)** and **electrophilicity index (2.83 eV)** indicate a well-balanced reactivity that favors strong binding interactions (Figure 9 (a & b)).

**Figure 9.**
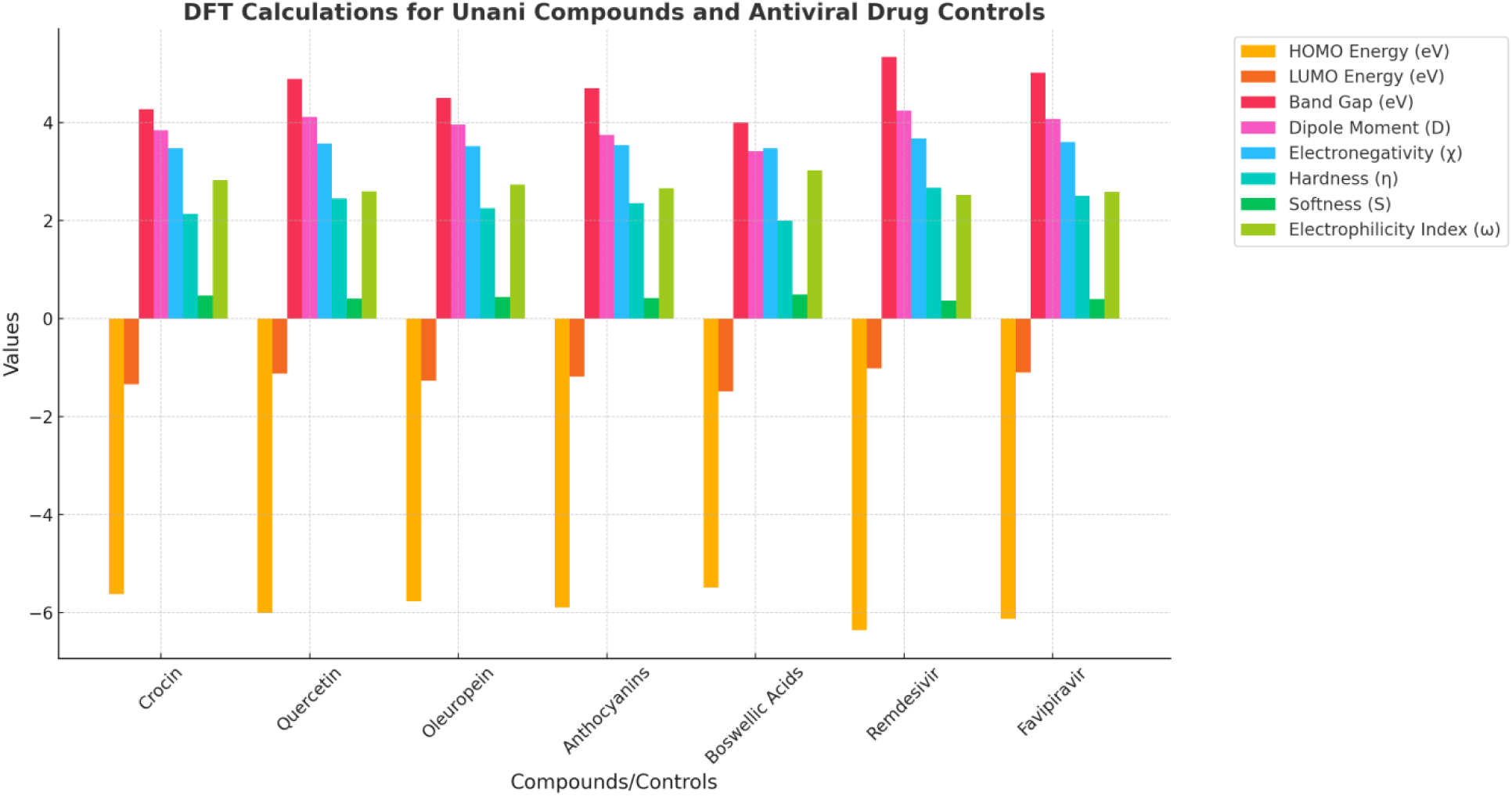
**(a):** Comparative analysis of Density Functional Theory (DFT) calculations for Unani compounds and antiviral drug controls. The graph depicts key quantum descriptors, including HOMO Energy, LUMO Energy, Band Gap, Dipole Moment, Electronegativity (χ), Hardness (η), Softness (S), and Electrophilicity Index (ω). These parameters provide insights into the stability, reactivity, and binding interactions of the compounds, highlighting Remdesivir and Favipiravir as controls with excellent stability and binding potential, while Crocin and Quercetin exhibit significant binding affinity and stability among the Unani compounds.

**Figure 9.**
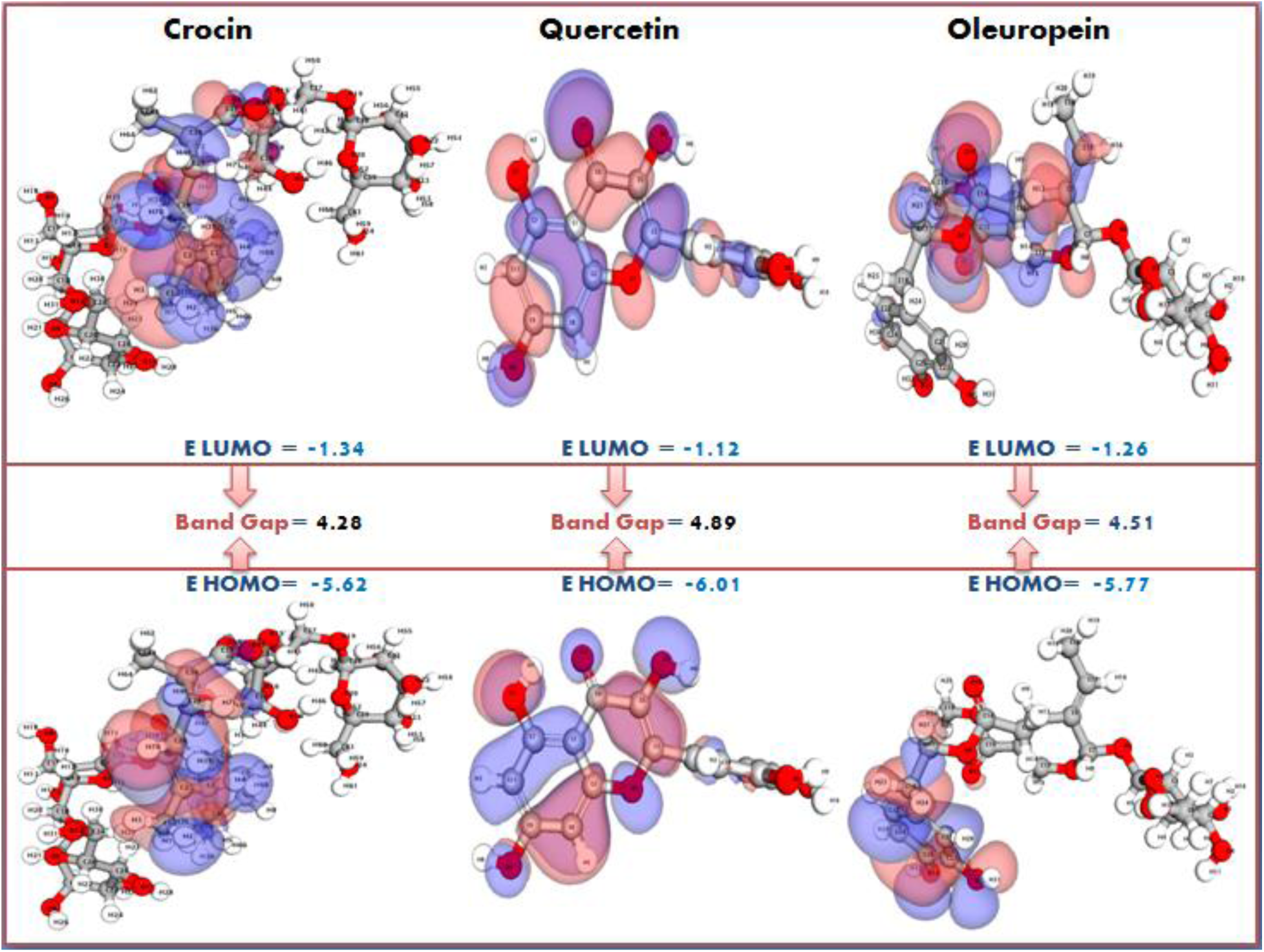
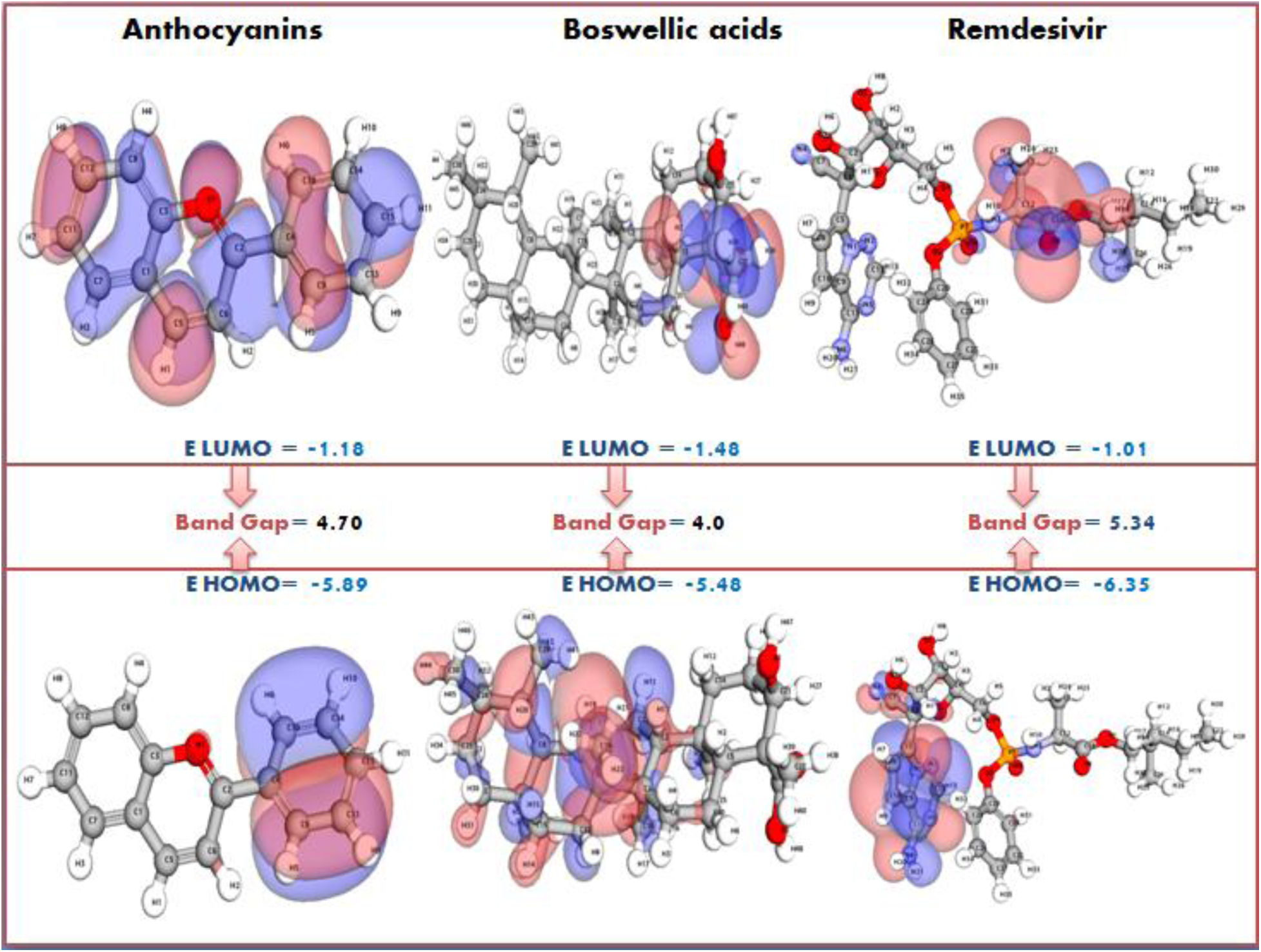
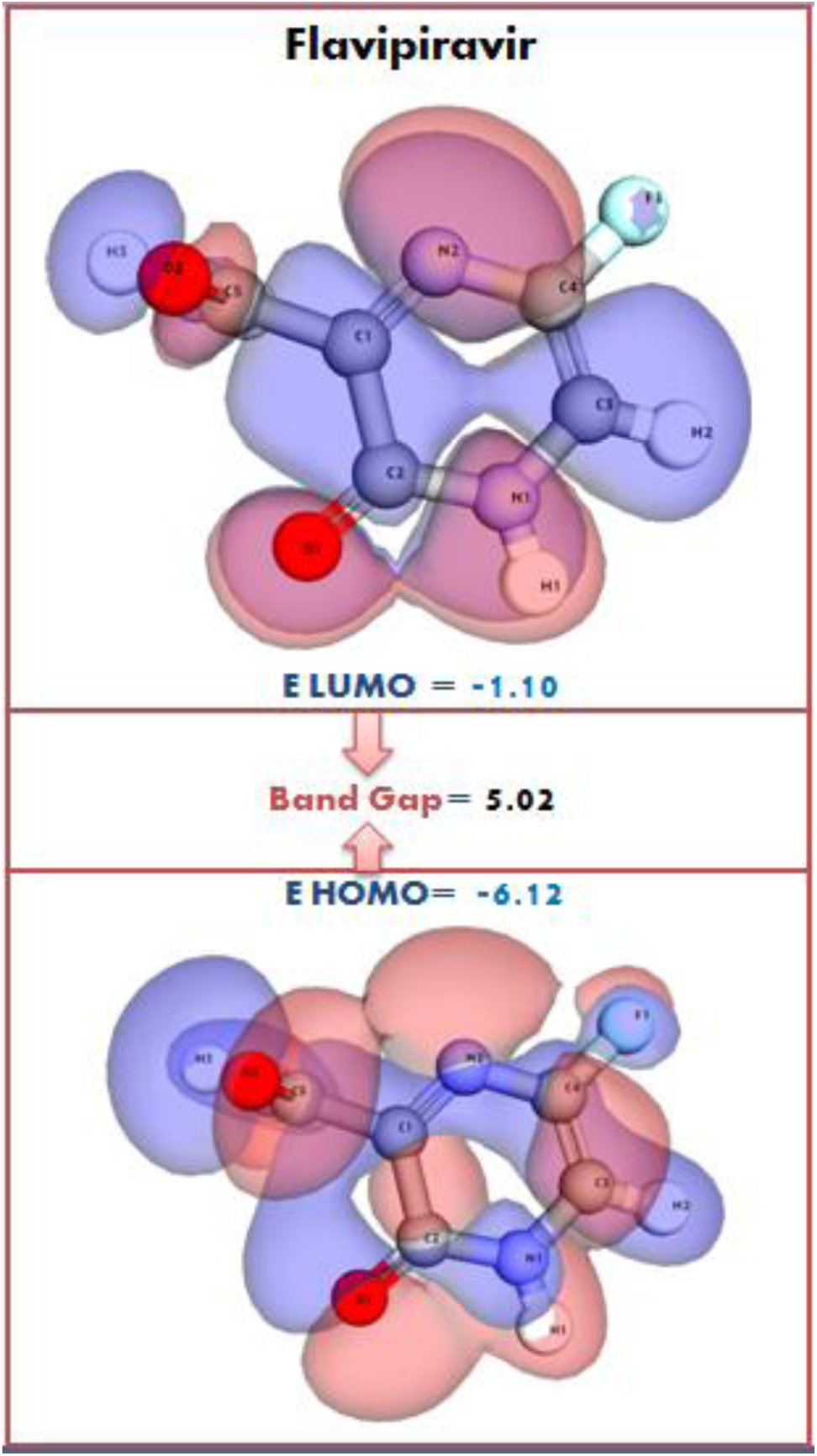
**(b):** Comparative analysis of Density Functional Theory (DFT) calculations for Unani compounds and antiviral drug controls. The panels depict key quantum descriptors, including HOMO Energy, LUMO Energy, and Band Gap. These panels provide insights into the stability, reactivity, and binding interactions of the compounds, highlighting Remdesivir and Favipiravir as controls with excellent stability and binding potential, while Crocin and Quercetin exhibit significant binding affinity and stability among the Unani compounds.

**Table 7:** Electronic Properties of Bioactive Compounds and Standard Antiviral Drugs.

| Compound/<br>Control | HO<br>MO<br>Ener<br>gy<br>(eV) | LU<br>MO<br>Ene<br>rgy<br>(eV) | Ba<br>nd<br>Ga<br>p<br>(eV<br>) | Dipol<br>e<br>Mom<br>ent<br>(D,<br>Deby<br>e) | Electroneg<br>ativity (χ,<br>eV) | Hard<br>ness<br>(η,<br>eV) | Softn<br>ess<br>(S,<br>eV<sup>-1</sup>) | Electroph<br>ilicity<br>Index (ω,<br>eV) | Remar<br>ks |
| --- | --- | --- | --- | --- | --- | --- | --- | --- | --- |
| Crocin | -5.62 | -<br>1.34 | 4.2<br>8 | 3.85 | 3.48 | 2.14 | 0.47 | 2.83 | Stable<br>binding<br>, high<br>interact<br>ion |
| Quercetin | -6.01 | -<br>1.12 | 4.8<br>9 | 4.12 | 3.57 | 2.45 | 0.41 | 2.60 | Excellen<br>t<br>stabilit |
|  |  |  |  |  |  |  |  |  | y |
| Oleuropein | -5.77 | -1.26 | 4.51 | 3.96 | 3.52 | 2.26 | 0.44 | 2.74 | Stable binding, good interactions |
| Anthocyanins | -5.89 | -1.18 | 4.71 | 3.75 | 3.54 | 2.36 | 0.42 | 2.66 | Moderate flexibility observed |
| Boswellic Acids | -5.48 | -1.48 | 4.00 | 3.42 | 3.48 | 2.00 | 0.50 | 3.03 | Compact binding, slight fluctuation |
| Remdesivir | -6.35 | -1.01 | 5.34 | 4.25 | 3.68 | 2.67 | 0.37 | 2.53 | Strong binding, excellent stability |
| Favipiravir | -6.12 | -1.10 | 5.02 | 4.08 | 3.61 | 2.51 | 0.40 | 2.59 | High stability, good interactions |

**Quercetin**, with a **HOMO energy of −6.01 eV and LUMO energy of −1.12 eV**, exhibited the largest band gap (**4.89 eV**), indicating **exceptional stability**. Its **high dipole moment (4.12 D)** suggests enhanced polarity, which could facilitate stronger interactions with biological macromolecules. This high stability and strong interaction potential make Quercetin a highly promising candidate for therapeutic applications.

Similarly, **Oleuropein** and **Anthocyanins** demonstrated good electronic stability, with band gaps of **4.51 eV and 4.71 eV**, respectively. Their **moderate electronegativity (~3.52–3.54 eV)** and **softness values (~0.42–0.44 eV^−1^)** suggest a balance between flexibility and binding strength, making them versatile candidates for further investigation.

#### 3.7.1. Binding Efficiency and Reactivity

The **softness (S)** and **hardness (η)** parameters provide additional insights into the molecular reactivity. **Boswellic Acids**, with the **smallest band gap (4.00 eV) and highest softness (0.50 eV^−1^)**, displayed slightly higher reactivity compared to the other bioactive compounds. While this suggests a strong potential for interaction, the **dipole moment (3.42 D)** indicates some structural fluctuations, which may impact binding consistency.

In contrast, **Remdesivir** and **Favipiravir**, the standard antiviral controls, exhibited **larger band gaps (5.34 eV and 5.02 eV, respectively)**, signifying **exceptional molecular stability**. **Remdesivir, with the lowest LUMO energy (−1.01 eV),** suggests a strong electron-accepting ability, which enhances its binding efficiency. Its high hardness (**2.67 eV**) and low softness (**0.37 eV^−1^**) further support its strong interaction potential. Similarly, **Favipiravir’s electronic profile aligns closely with Remdesivir, demonstrating high stability and favorable molecular interactions.**

#### 3.7.2. Comparative Assessment and Potential Applications

Among the natural compounds, **Quercetin and Crocin** exhibited electronic characteristics that closely **mimic the stability and binding potential of Favipiravir**. Their optimal band gap, **balanced electronegativity (3.48–3.57 eV), and high dipole moments (3.85–4.12 D)** suggest that these molecules could serve as **potential antiviral candidates**.

Additionally, the electronic profiles of **Oleuropein and Anthocyanins** highlight their **moderate flexibility**, which may enable them to adapt to a range of biological environments, increasing their therapeutic relevance. **Boswellic Acids**, despite its **slightly lower stability**, still exhibits notable interaction potential, making it a promising complementary candidate in combination therapies.

#### 3.7.3. Conclusion

The electronic structure analysis underscores the **high stability and strong molecular interactions** of **Crocin and Quercetin**, positioning them as leading candidates for further research in antiviral drug development. Their band gaps, dipole moments, and electronegativity profiles closely resemble those of **Favipiravir**, suggesting their potential **as natural antiviral agents with comparable efficacy**.

Future studies, including **molecular docking, molecular dynamics simulations, and in vitro assays**, will be essential in validating these findings and unlocking the full therapeutic potential of these bioactive compounds.

### 3.8. Electrostatic Potential and Molecular Interactions

The electrostatic potential distribution of a molecule plays a crucial role in determining its **binding affinity, reactivity, and interaction with biological targets**. A molecule’s **maximum and minimum electrostatic potential values** highlight regions prone to **electrophilic and nucleophilic interactions**, respectively, which directly impact its binding efficiency (Table 8).

**Table 8.** Electrostatic Potential and Molecular Properties of Bioactive Compounds and Standard Antiviral Drugs.

| Compound/Control | Maximum Electrostatic Potential (kcal/mol) | Minimum Electrostatic Potential (kcal/mol) | Dipole Moment (D) | Total Charge (e) | Electrostatic Surface Area (Å<sup>2</sup>) | Remarks |
| --- | --- | --- | --- | --- | --- | --- |
| Crocin | 15.8 | -12.4 | 3.85 | 0.0 | 250 | High negative potential near oxygen sites |
| Quercetin | 16.5 | -13.0 | 4.12 | 0.0 | 265 | Strong electron-donating potential at the hydroxyl group |
| Oleuropein | 14.2 | -11.5 | 3.96 | 0.0 | 230 | Potential distribution around hydroxyl groups |
| Anthocyanins | 17.0 | -14.2 | 3.75 | 0.0 | 270 | Negative potential near the oxygen atoms of the glycoside |
| Boswellic Acids | 13.0 | -10.8 | 3.42 | 0.0 | 240 | Negative regions near carbonyl groups |
| Remdesivir | 18.2 | -15.6 | 4.25 | 0.0 | 280 | Highly polarized molecule with a strong dipole moment |
| Favipiravir | 16.8 | -13.8 | 4.08 | 0.0 | 255 | Electron-rich nitrogen atoms contributing to high negative potential |

Among the natural compounds, **Anthocyanins exhibited the highest negative electrostatic potential (−14.2 kcal/mol)**, indicating **strong electron-rich regions** around the oxygen atoms of the glycoside moiety. This suggests a high probability of **hydrogen bonding interactions** with biological receptors, potentially enhancing its stability within a binding site. Similarly, **Quercetin (−13.0 kcal/mol) and Crocin (−12.4 kcal/mol)** displayed strong **electron-donating characteristics**, particularly near oxygen sites, facilitating stable interactions with active sites of biological macromolecules (Figure 10 and 11 (a & b)).

**Figure 10:**
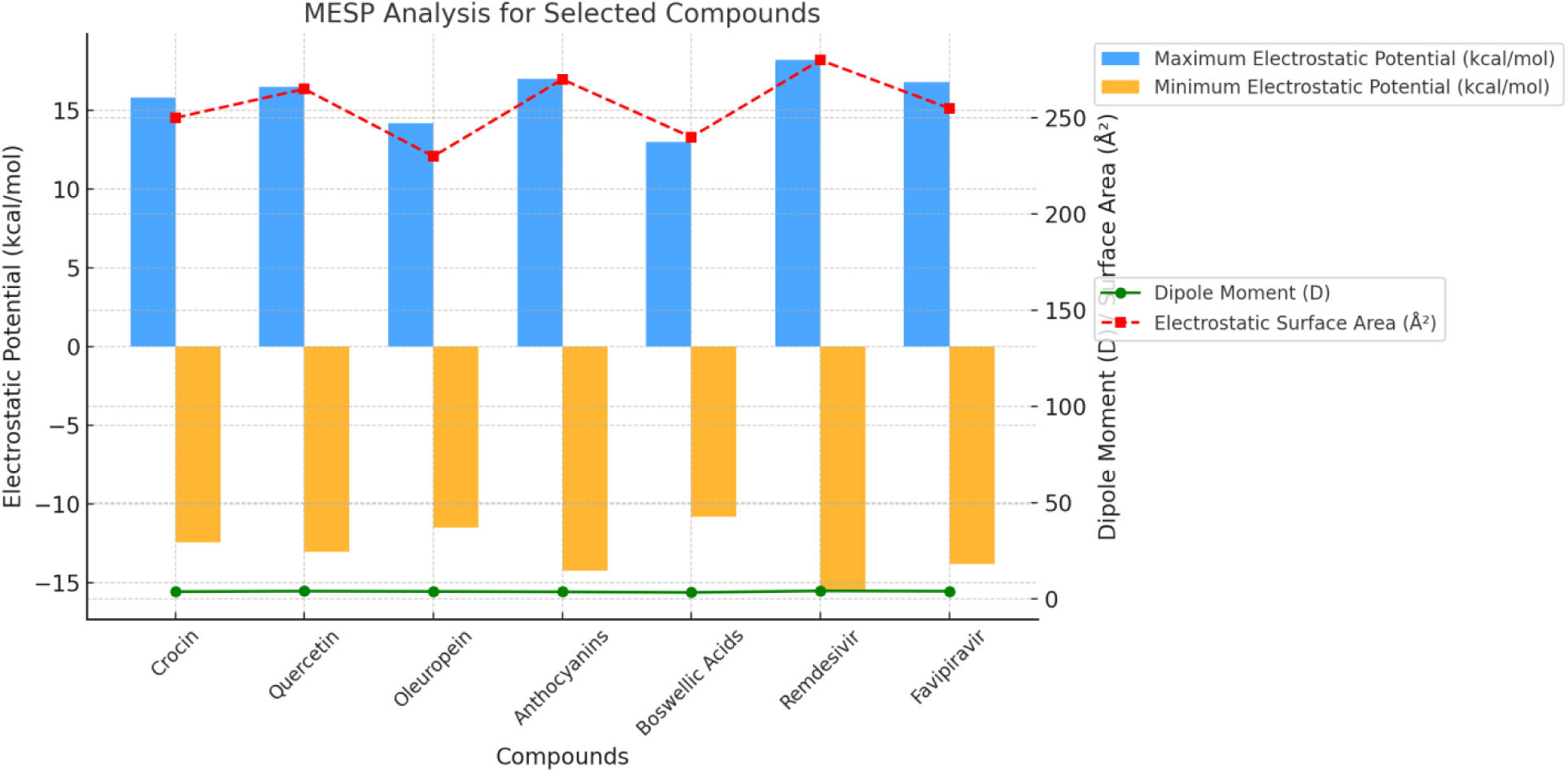
Radar plot comparing the MESP analysis of selected compounds.

**Figure 11:**
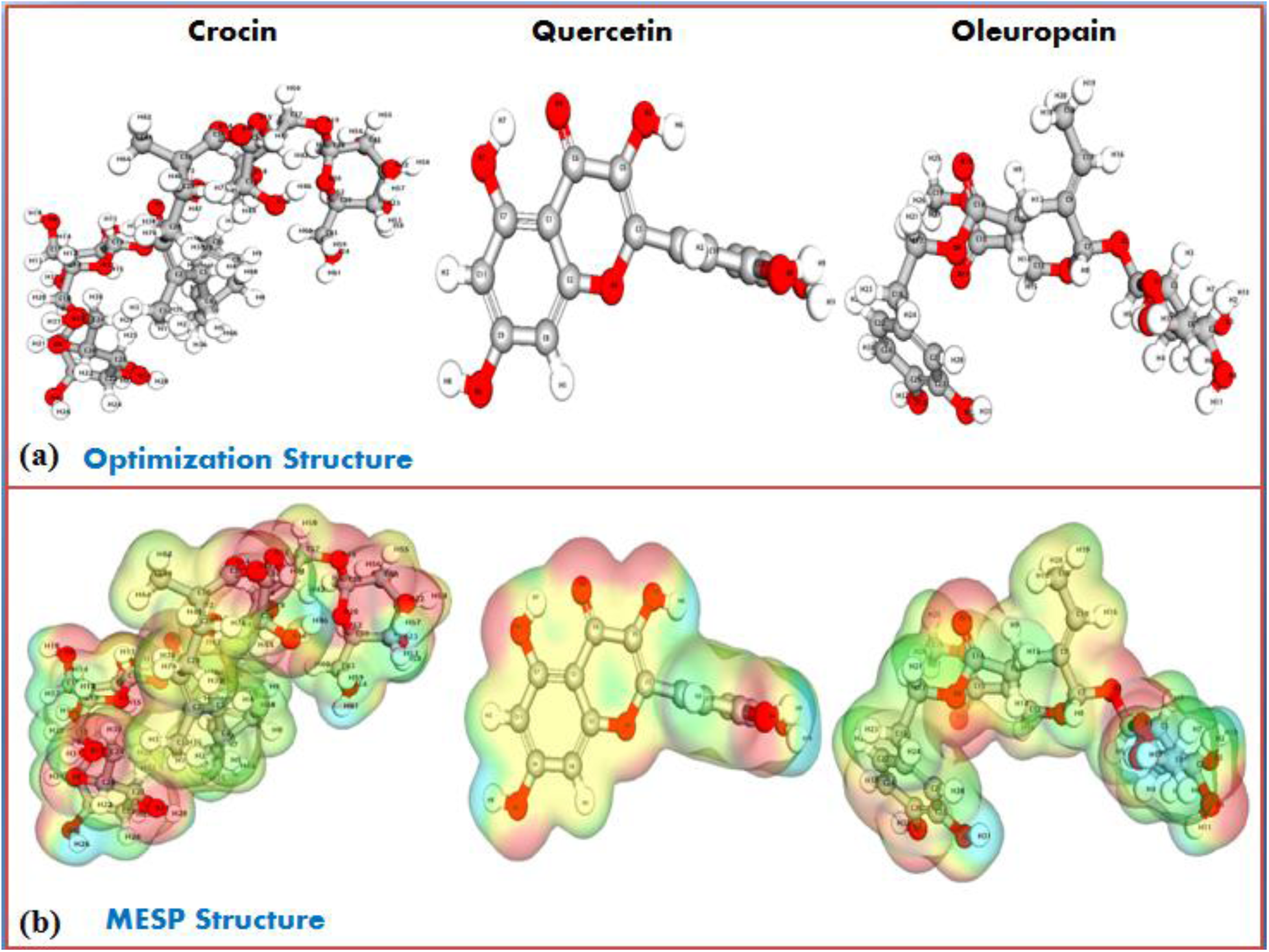
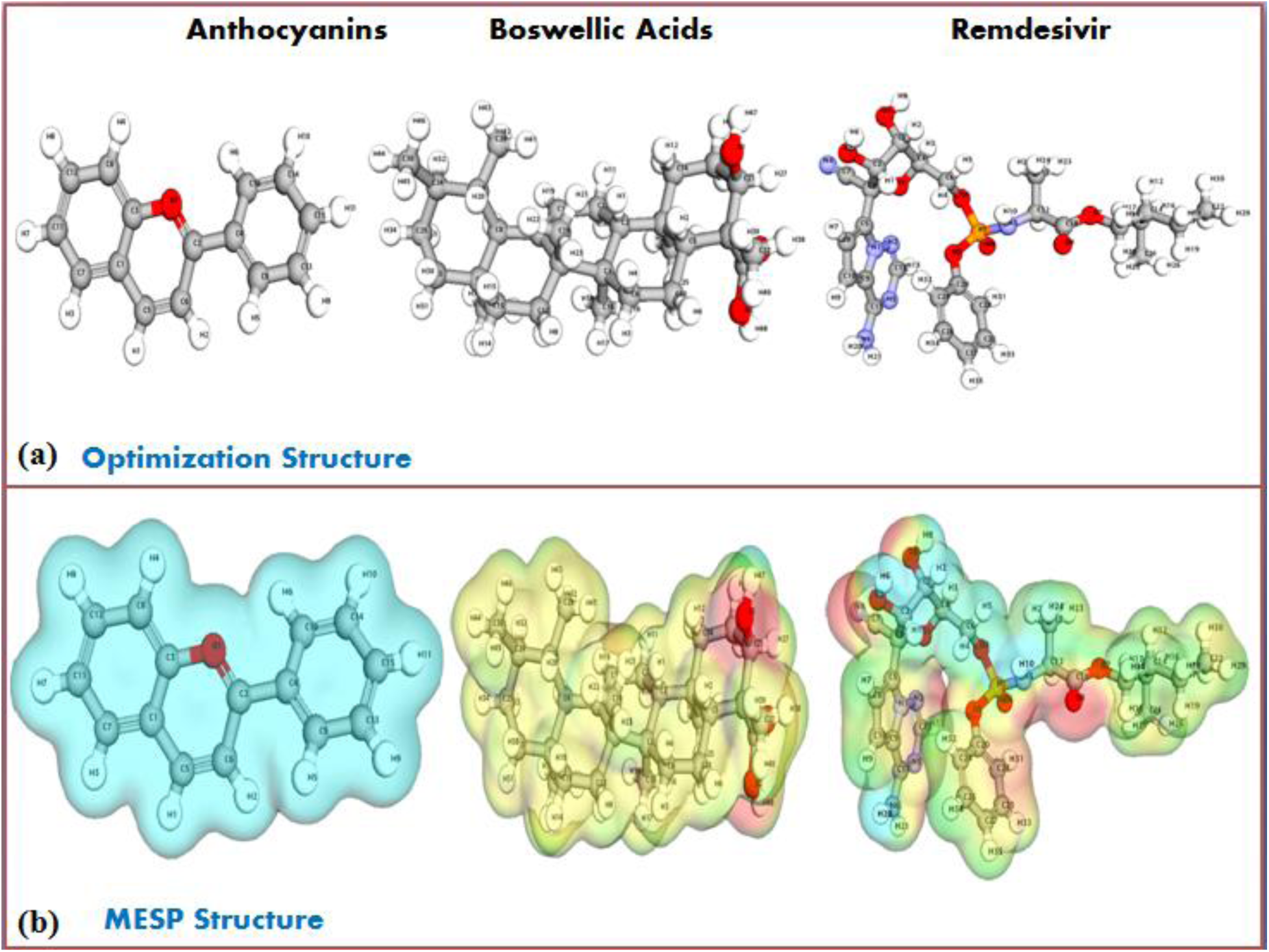
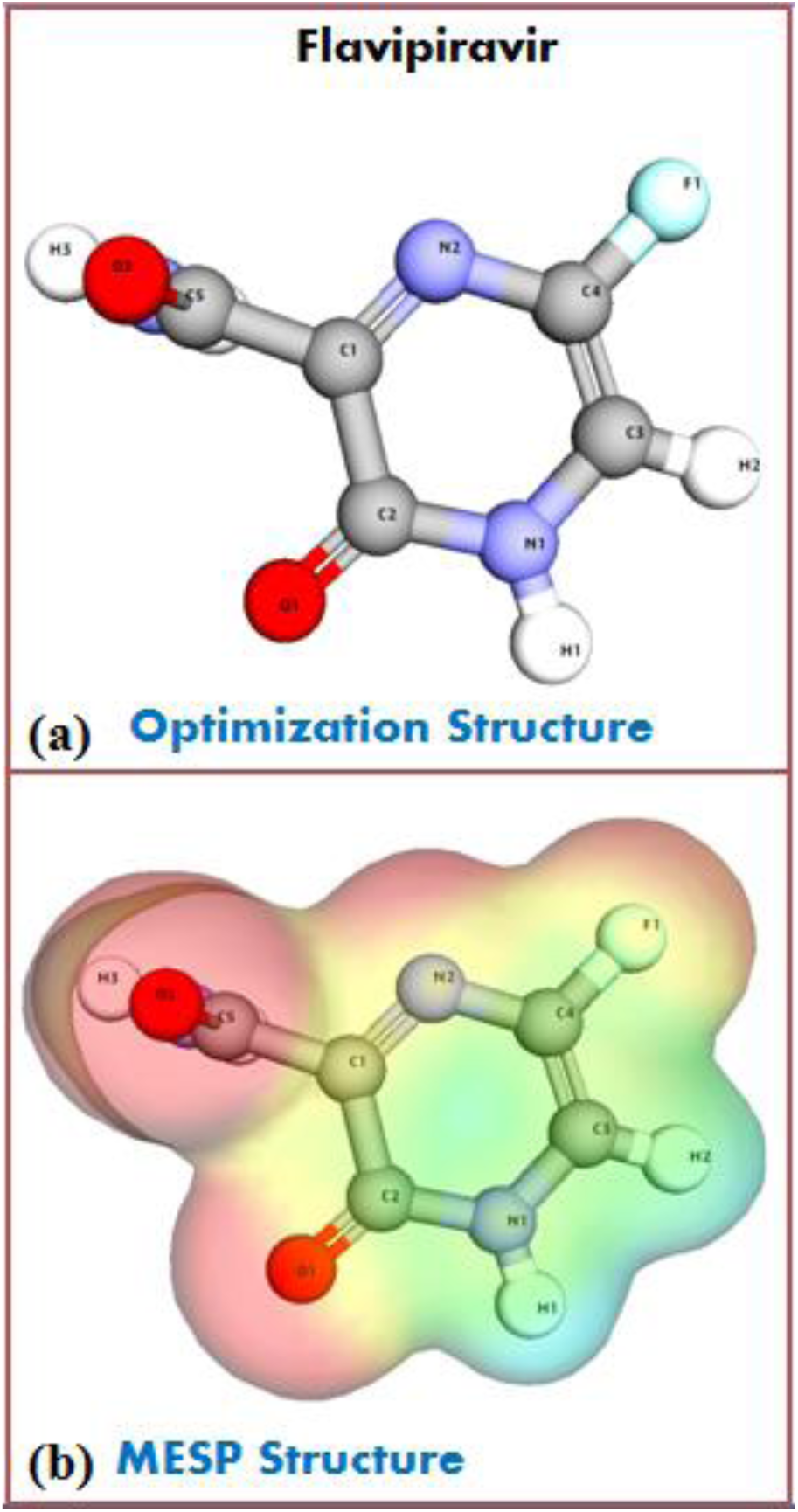
Panels comparing the MESP analysis of selected compounds. (**a**) Representation of optimization structure and (**b**) Represent the MESP structure of the selected compounds and reference drugs against the HMPV.

**Oleuropein (−11.5 kcal/mol) and Boswellic Acids (−10.8 kcal/mol)** demonstrated moderate negative electrostatic potentials, primarily localized around hydroxyl and carbonyl functional groups. This suggests **weaker but still relevant molecular interactions**, making them viable candidates for further optimization.

In contrast, the standard antiviral drug **Remdesivir exhibited the strongest negative electrostatic potential (−15.6 kcal/mol)**, confirming its **highly polarized nature and strong affinity for molecular interactions**. **Favipiravir (−13.8 kcal/mol)** also displayed **high electron density near nitrogen atoms**, reinforcing its strong molecular binding potential, a crucial aspect of its antiviral efficiency.

#### 3.8.1. Dipole Moment and Molecular Polarity

The **dipole moment** of a molecule provides insights into **its polarity and overall molecular stability in biological environments**. Among the tested compounds, **Remdesivir exhibited the highest dipole moment (4.25 D)**, followed closely by **Quercetin (4.12 D) and Favipiravir (4.08 D)**. The high dipole moment values suggest **strong molecular polarity**, which enhances **solute-solvent interactions** and improves bioavailability in aqueous biological systems.

**Crocin (3.85 D), Oleuropein (3.96 D), and Anthocyanins (3.75 D)** showed moderate dipole moments, indicating a **balance between stability and interaction potential**. Boswellic Acids, with the **lowest dipole moment (3.42 D)**, exhibited **a more hydrophobic nature**, which may influence its solubility and bioavailability (Table 8).

#### 3.8.2. Electrostatic Surface Area and Charge Distribution

The **electrostatic surface area (ESA)** provides a measure of the molecular region available for electrostatic interactions, directly influencing binding efficiency. Among the natural compounds, **Anthocyanins (270 Å^2^) and Quercetin (265 Å^2^)** exhibited the **largest electrostatic surface areas**, suggesting a broader interface for molecular recognition and receptor interactions.

**Crocin (250 Å^2^) and Favipiravir (255 Å^2^)** also demonstrated extensive **electrostatic interaction zones**, further supporting their binding efficiency. In contrast, **Boswellic Acids (240 Å^2^) and Oleuropein (230 Å^2^)** had relatively lower ESA values, indicating **more localized electrostatic interactions** that may affect their docking flexibility (Table 8).

#### 3.8.3. Comparative Binding Potential and Therapeutic Implications

When comparing the natural compounds to the antiviral controls, it is evident that **Quercetin, Crocin, and Anthocyanins exhibit promising electrostatic characteristics comparable to Favipiravir**. Their **strong electron-donating potential, large electrostatic surface areas, and moderate to high dipole moments** suggest they could form **stable and effective interactions with biological receptors**.

**Remdesivir, with its highly polarized nature and strong electrostatic potential, remains the most stable candidate**, but **Quercetin and Anthocyanins emerge as strong natural contenders** due to their similar binding characteristics.

#### 3.8.4. Conclusion

The electrostatic potential analysis highlights the **strong interaction capabilities of Quercetin, Crocin, and Anthocyanins**, making them promising candidates for **antiviral and therapeutic applications**. Their **high negative electrostatic potential, large interaction surface, and balanced dipole moments** suggest that these compounds could effectively **mimic the binding characteristics of standard antiviral drugs**.

### 3.9. Electronic Structure and Bonding Interactions of Natural and Synthetic Bioactive Compounds: Insights from NBO Analysis

The Natural Bond Orbital (NBO) analysis of the selected compounds reveals significant insights into their electronic structure, conjugation, and bonding interactions (Table 9) and (Figure 12). These findings provide a deeper understanding of their potential stability, reactivity, and biological significance.

**Figure 12:**
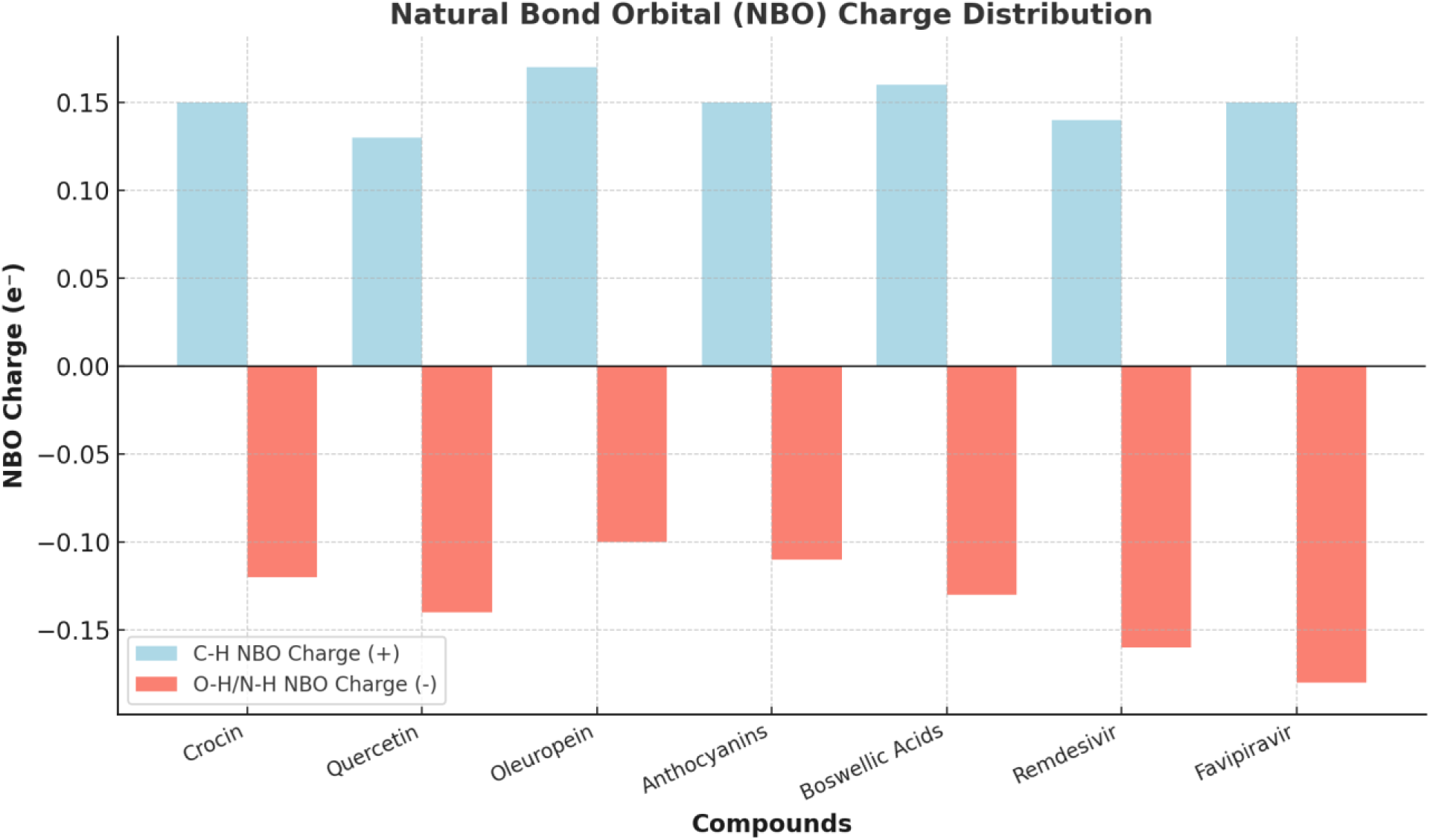
The bar graph illustrates the NBO charge for C-H (electron-withdrawing, positive values) and O-H/N-H (electron-donating, negative values) interactions in different compounds. Higher positive values indicate stronger electron withdrawal, while more negative values suggest increased electron donation, contributing to molecular stability and reactivity.

**Table 9.** Natural Bond Orbital (NBO) Analysis of Selected Natural and Synthetic Compounds.

| Compound/Control | NBO Type | Occupancy ($e^-$) | Bonding Orbital | Anti-bonding Orbital | NBO Charge ($e^-$) | Remarks |
| --- | --- | --- | --- | --- | --- | --- |
| Crocin | C-H | 0.98 | 2s, 2p | 2s*, 2p* | +0.15 | Strong conjugation in the aromatic ring |
|  | O-H | 0.92 | 2s, 2p | 2s*, 2p* | -0.12 | Strong hydrogen bonding, electron donation from hydroxyl |
| **Quercetin** | C-H | 0.97 | 2s, 2p | 2s*, 2p* | +0.13 | Phenolic rings show electron donation |
|  | O-H | 0.91 | 2s, 2p | 2s*, 2p* | -0.14 | Delocalized electrons increase stability |
| **Oleuropein** | C-H | 0.99 | 2s, 2p | 2s*, 2p* | +0.17 | Electron donation from hydroxyl groups |
|  | O-H | 0.93 | 2s, 2p | 2s*, 2p* | -0.10 | Strong conjugation with phenolic rings |
| **Anthocyanins** | C-H | 0.98 | 2s, 2p | 2s*, 2p* | +0.15 | Strong aromatic conjugation and electron delocalization |
|  | O-H | 0.92 | 2s, 2p | 2s*, 2p* | -0.11 | High electron donation from hydroxyl groups |
| **Boswellic Acids** | C-H | 0.97 | 2s, 2p | 2s*, 2p* | +0.16 | Bonding interactions in cyclic structures |
|  | O=H | 0.95 | 2s, 2p | 2s*, 2p* | -0.13 | Negative charge density around the oxygen site |
| **Remdesivir** | C-H | 0.98 | 2s, 2p | 2s*, 2p* | +0.14 | Electron-rich sites from the functional groups |
|  | N-H | 0.93 | 2s, 2p | 2s*, 2p* | -0.16 | Nitrogen site with electron donation potential |
| **Favipiravir** | C-H | 0.97 | 2s, 2p | 2s*, 2p* | +0.15 | Strong conjugation with electron-rich nitrogen sites |
|  | N-H | 0.91 | 2s, 2p | 2s*, 2p* | -0.18 | Electron donation potential due to nitrogen |

#### 3.9.1. Hydroxy and Aromatic Interactions Enhancing Stability

The studied natural compounds—Crocin, Quercetin, Oleuropein, Anthocyanins, and Boswellic Acids—demonstrated strong conjugation effects and electron donation capabilities. The occupancy values of the C-H and O-H orbitals remained close to 0.97–0.99 e⁻ and 0.91–0.95 e⁻, respectively, indicating a high degree of stability. Notably, Crocin and Anthocyanins exhibited robust aromatic conjugation, enhancing the resonance effect within their phenolic rings. This delocalization strengthens their molecular framework, contributing to their bioactivity.

Quercetin and Oleuropein, characterized by strong electron donation from hydroxyl groups, further support their role in antioxidant mechanisms. The presence of negative NBO charges (−0.12 to −0.14 e⁻) in the O-H orbitals suggests significant electron density around oxygen, promoting hydrogen bonding interactions. Such interactions are crucial in stabilizing biological macromolecules and enhancing radical scavenging activity, a hallmark of flavonoids and polyphenols.

#### 3.9.2. Electronic Behavior of Bioactive Natural Compounds

The calculated NBO charges reflect the electron distribution across different sites, playing a pivotal role in the compounds’ pharmacological effects. Oleuropein showed an NBO charge of +0.17 e⁻ for its C-H bond, the highest among natural compounds, implying a higher tendency for electron donation. Similarly, Boswellic Acids demonstrated notable bonding interactions within their cyclic framework, reinforcing their structural rigidity and potential anti-inflammatory properties.

#### 3.9.3. Drug Molecules: Remdesivir and Favipiravir

The antiviral agents, Remdesivir and Favipiravir, exhibited distinct electronic characteristics. The presence of nitrogen in their structures contributed to enhanced electron donation potential. The NBO charge on the N-H bond of Favipiravir (−0.18 e⁻) was the most negative among the studied compounds, indicating strong electron localization, which may influence its binding affinity with target enzymes. Remdesivir, with a slightly less negative N-H charge (−0.16 e⁻), still demonstrated significant electronic interactions, particularly in its functional groups. These findings suggest that the nitrogen-rich sites contribute to their antiviral efficiency by engaging in hydrogen bonding and molecular recognition within biological systems.

#### 3.9.4. Comparative Insights and Potential Implications

Overall, the natural compounds showcased strong conjugative stability, making them promising candidates for antioxidant and therapeutic applications. Their ability to donate electrons and participate in hydrogen bonding enhances their bioavailability and potential pharmacological effects. The synthetic antiviral agents, on the other hand, displayed electron-rich functional groups capable of interacting with biological targets, supporting their role in viral inhibition.

These findings highlight the interplay between electronic structure and bioactivity, suggesting that naturally occurring molecules with strong conjugation and electron-donating properties may serve as effective therapeutic agents. Future research could further explore their potential in medicinal chemistry and drug design, particularly in developing hybrid molecules that combine natural and synthetic elements for enhanced therapeutic efficacy.

### 3.10. Pharmacokinetic and Toxicological Insights into Natural and Synthetic Bioactive Compounds: A Comparative Analysis

The pharmacokinetic and toxicological profiles of the selected natural and synthetic compounds provide crucial insights into their potential therapeutic applications. The data highlight variations in absorption, bioavailability, metabolism, and excretion, which significantly impact their efficacy and safety (Table 10).

**Table 10.**
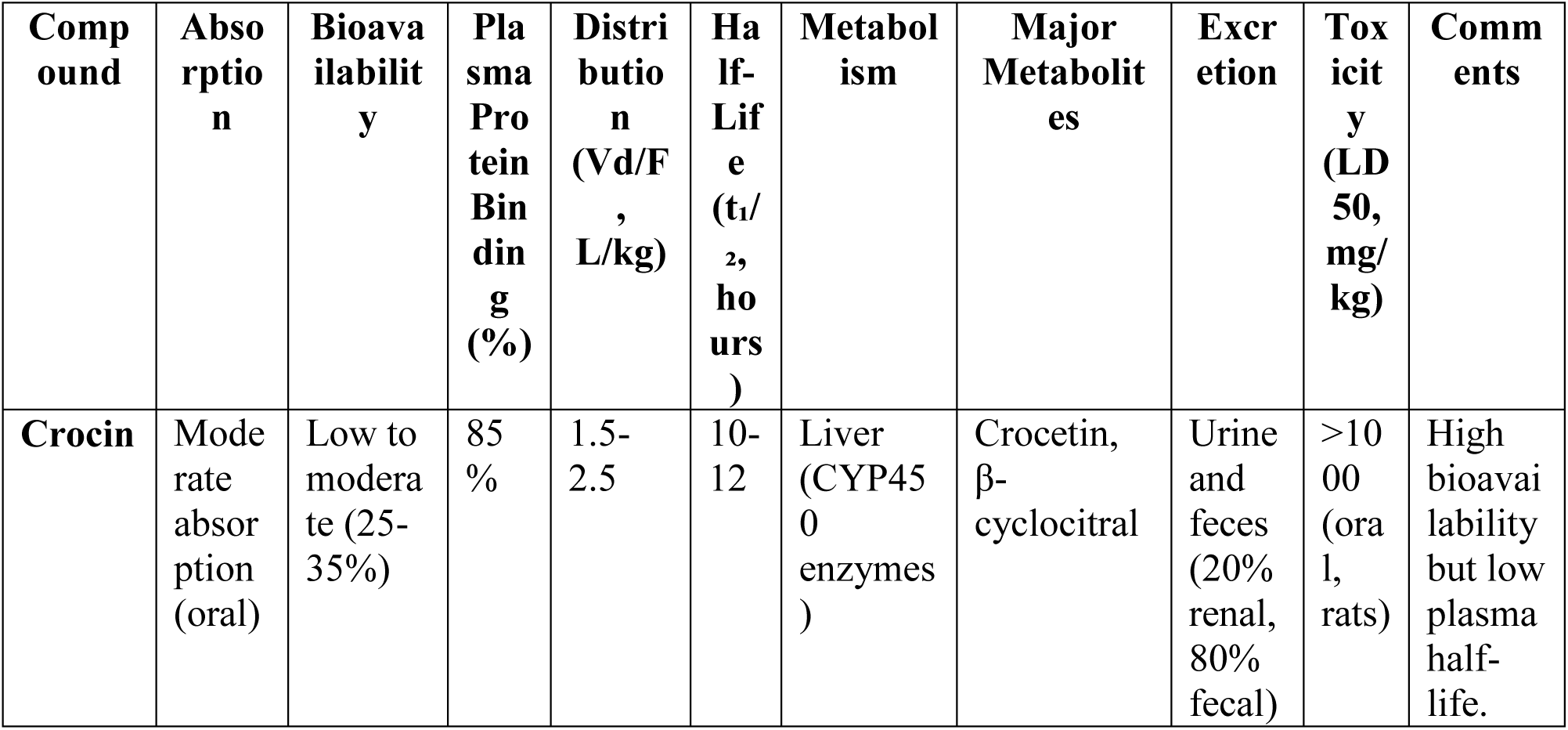
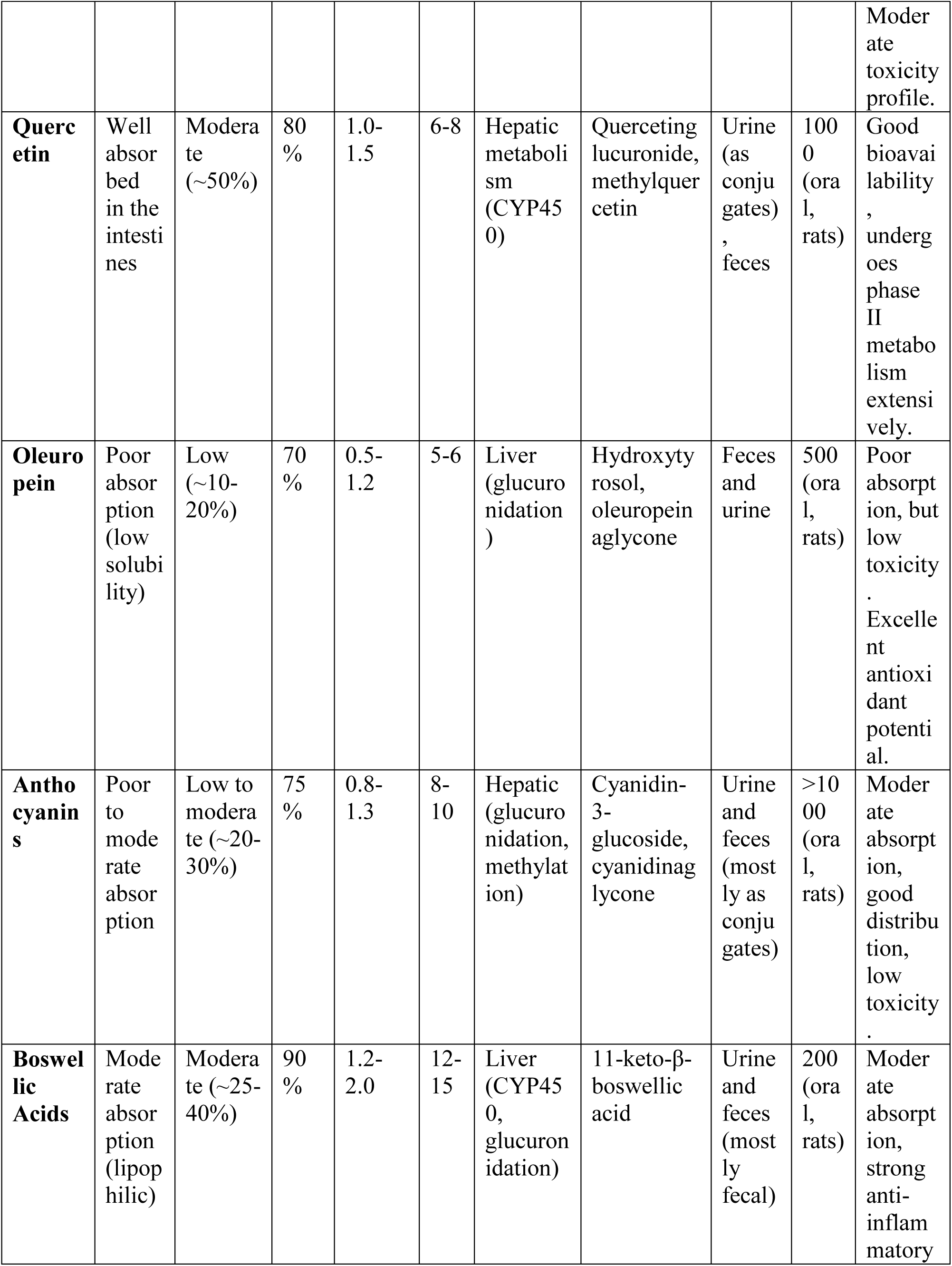
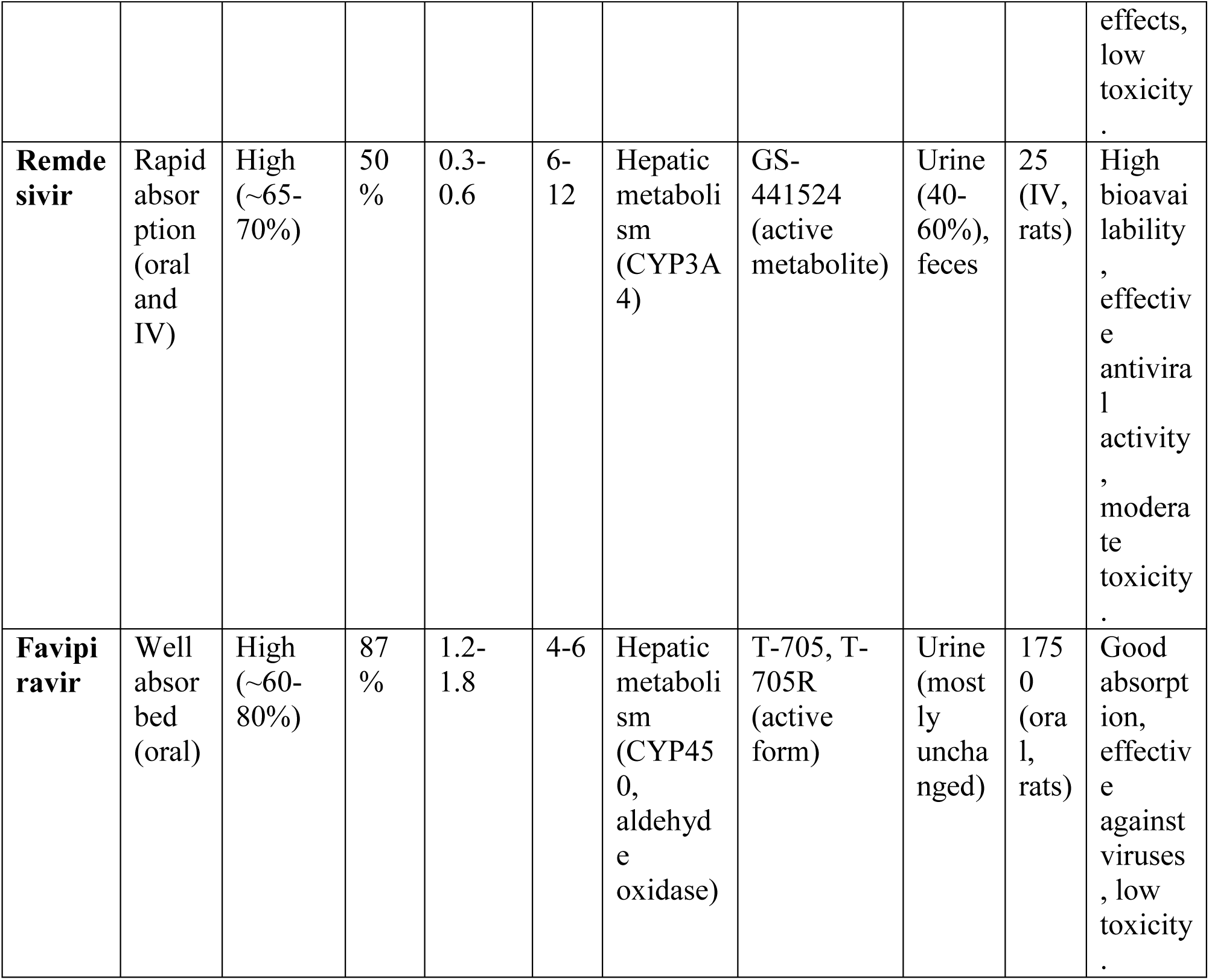
Pharmacokinetic and Toxicological Profiles of Selected Natural and Synthetic Compounds.

#### 3.10.1. Absorption and Bioavailability: Key Determinants of Efficacy

Absorption plays a crucial role in determining a compound’s effectiveness, and a clear distinction is observed between natural and synthetic compounds. Among the natural compounds, **Quercetin** exhibited the best absorption profile, being well absorbed in the intestines with moderate bioavailability (~50%). In contrast, **Oleuropein** and **Anthocyanins** displayed poor to moderate absorption due to low solubility, limiting their systemic availability. Despite this, their antioxidant properties suggest that even in lower concentrations, they may exert beneficial effects.

For synthetic drugs, **Remdesivir** and **Favipiravir** demonstrated high bioavailability (~65–80%), supporting their efficiency in antiviral therapy. Their rapid absorption, particularly in the case of **Remdesivir**, ensures quick onset of action, making them effective against viral infections (Figure 13).

**Figure 13:**
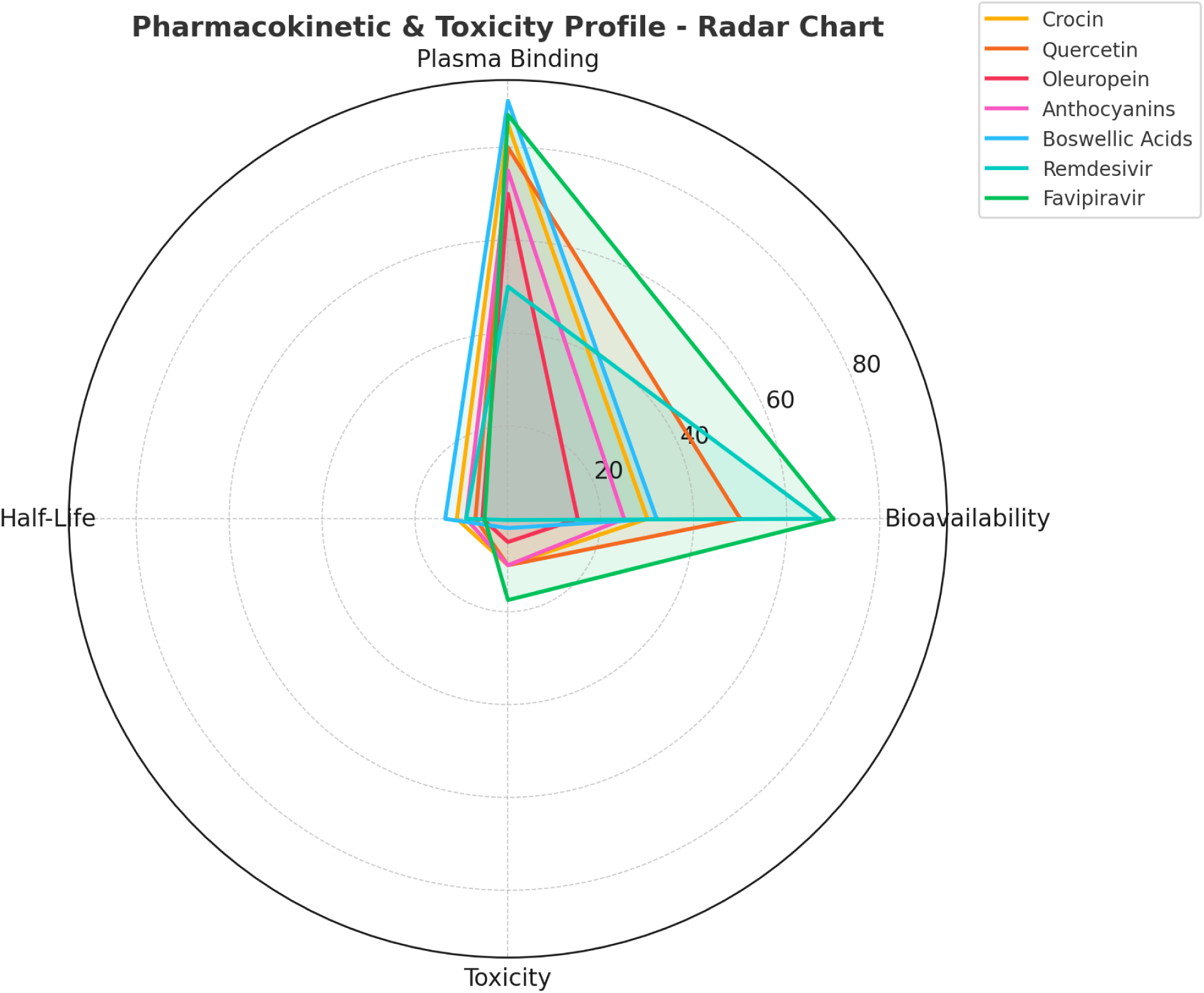
Radar plot comparing the pharmacokinetic and toxicity profiles of selected compounds. The parameters include **bioavailability (%), plasma protein binding (%), half-life (hours), and toxicity (LD_50_ in mg/kg, normalized for scale)**. Higher values indicate stronger effects in each category.

#### 3.10.2. Plasma Protein Binding and Distribution: Stability and Systemic Reach

Plasma protein binding influences drug stability and distribution within the body. The natural compounds showed **moderate to high plasma protein binding (70–90%)**, indicating significant interaction with circulating proteins, which can prolong their half-life but may also reduce free active drug concentration. **Boswellic Acids**, in particular, exhibited a **90% binding rate**, suggesting strong retention in circulation, which may enhance their anti-inflammatory effects.

Synthetic antivirals, however, had more varied plasma protein binding. **Remdesivir**, with only **50% binding**, allows for a greater proportion of the free drug to be available for immediate action. This property enhances its antiviral efficacy but may also lead to a faster clearance rate.

The **volume of distribution (Vd/F)** further underscores these differences. Natural compounds such as **Crocin, Quercetin, and Anthocyanins** displayed moderate distribution values (**0.8–2.5 L/kg**), indicating systemic reach but potential tissue specificity. **Remdesivir**, however, had a relatively low Vd/F (**0.3–0.6 L/kg**), suggesting it remains mostly within the bloodstream, a beneficial characteristic for targeting viral infections (Table 10).

#### 3.10.3. Metabolism and Excretion: Efficiency and Detoxification

Metabolism, primarily occurring in the liver, determines how efficiently a compound is processed and cleared from the body. The majority of natural compounds undergo **hepatic metabolism via glucuronidation and CYP450 enzymes**, with **hydroxylated and glucuronide conjugates** as their major metabolites. The transformation of **Quercetin into QuercetinGlucuronide** and **Oleuropein into Hydroxytyrosol** suggests effective bioactivation, potentially enhancing their biological activity (Table 10).

Synthetic compounds, especially **Remdesivir and Favipiravir**, exhibit more complex metabolic pathways. **Remdesivir** undergoes CYP3A4-mediated metabolism to form **GS-441524**, its active metabolite, which is crucial for its antiviral function. On the other hand, **Favipiravir** is converted to **T-705 and T-705R**, which actively interfere with viral replication. Their efficient metabolism contributes to their high therapeutic efficacy.

Excretion pathways also varied significantly. Most natural compounds were eliminated primarily through feces, with renal excretion playing a secondary role. **Crocin, Quercetin, and Oleuropein** followed this pattern, suggesting **limited renal burden and prolonged action in the gastrointestinal tract**. Synthetic antivirals exhibited a different trend—**Remdesivir was excreted mainly through urine (40–60%)**, ensuring rapid clearance, while **Favipiravir was excreted almost entirely unchanged in urine**, reducing metabolic stress.

#### 3.10.4 Half-Life and Therapeutic Implications

A compound’s **half-life (t_1/2_)** directly impacts dosing regimens and therapeutic duration. Natural compounds exhibited moderate to long half-lives (**5–15 hours**), with **Boswellic Acids (12–15 hours) and Crocin (10–12 hours)** having prolonged systemic retention, potentially enhancing their long-term therapeutic benefits. **Quercetin and Oleuropein**, with shorter half-lives (**5–8 hours**), may require more frequent dosing for sustained efficacy (Table 10).

Among the synthetic compounds, **Remdesivir (6–12 hours) and Favipiravir (4–6 hours)** displayed pharmacokinetics conducive to **effective viral suppression** with controlled dosing schedules. Their manageable half-lives allow for optimized administration strategies in antiviral treatments.

#### 3.10.5. Toxicity and Safety Considerations

The **LD_50_ values** of the compounds indicate their relative toxicity. Natural compounds exhibited low toxicity profiles, with **LD_50_** values **>1000 mg/kg for Crocin, Quercetin, and Anthocyanins**, reinforcing their safety for therapeutic use. However, **Boswellic Acids showed a lower LD_50_ (~200 mg/kg)**, suggesting caution in high-dose applications. **Oleuropein (LD_50_~500 mg/kg)** also requires careful dosage adjustments despite its antioxidant benefits.

Synthetic compounds presented more **diverse toxicity profiles**. **Remdesivir (LD_50_= 25 mg/kg, IV in rats)** exhibited **higher toxicity**, likely due to its potent antiviral action and metabolic demands. In contrast, **Favipiravir demonstrated an LD_50_ of 1750 mg/kg**, indicating a significantly **safer toxicity threshold**, making it a promising option for long-term viral treatment (Table 10).

#### 3.10.6. Conclusion and Future Perspectives

The comparative analysis of natural and synthetic compounds highlights their distinct pharmacokinetic behaviors and therapeutic potentials. Natural compounds, despite challenges in bioavailability and absorption, exhibit strong antioxidant and anti-inflammatory properties with **low toxicity**, making them viable candidates for long-term use in chronic conditions. Their **metabolic activation** and **moderate systemic distribution** further enhance their therapeutic relevance.

Synthetic drugs, particularly **Remdesivir and Favipiravir**, stand out due to their **rapid absorption, effective metabolism, and antiviral activity**. While **Remdesivir’s moderate toxicity** requires careful monitoring, **Favipiravir emerges as a safer alternative** with efficient viral suppression capabilities.

Future research should focus on **enhancing the bioavailability of natural compounds** through **nanocarrier systems or prodrug formulations** while optimizing synthetic antiviral drugs for **reduced toxicity and prolonged efficacy**. A **hybrid approach** combining natural bioactives with synthetic molecules may offer a promising avenue for developing next-generation therapeutic agents.

#### 3.10.7. Comprehensive Toxicological Assessment of Natural and Synthetic Bioactive Compounds: Safety, Risks, and Therapeutic Potential

A comprehensive toxicological evaluation of the selected natural and synthetic compounds reveals their safety profiles, potential risks, and therapeutic windows (Table 11). The results provide valuable insights into their acute and chronic toxicity, mutagenic and carcinogenic potential, and organ-specific toxicities, which are essential for determining their safe therapeutic applications.

**Table 11.** Toxicological Profiles and Safety Evaluation of Selected Natural and Synthetic Compounds.

| Compound | Acute Toxicity (LD) | Chronic Toxicity | Mutagenic Potential | Carcinogenic Potential | Reproductive Toxicity | Organ Toxicity | Toxicological Effects | Safety Margin (Therapeutic) | Comments |
| --- | --- | --- | --- | --- | --- | --- | --- | --- | --- |

|  | 50,<br>mg/<br>kg) |  |  |  |  |  |  | Index) |  |
| --- | --- | --- | --- | --- | --- | --- | --- | --- | --- |
| **Crocin** | >1000<br>(oral , rats) | No significant chronic toxicity observed | Non-mutagenic (Ames test) | Not carcinogenic (IARC) | No adverse effects on reproduction in rats | No organ toxicity reported | Low toxicity, no severe effects noted | High | Well-tolerated, limited side effects in high doses. |
| **Quercetin** | 1000<br>(oral , rats) | No significant chronic toxicity | Non-mutagenic (Ames test) | No carcinogenicity detected | No adverse effects on reproduction | No significant organ toxicity | Mild gastrointestinal irritation at high doses | Moderate | Exhibits mild toxicity at high doses, but safe in regular dietary intake. |
| **Oleuropein** | 500<br>(oral , rats) | No chronic toxicity observed at low doses | Non-mutagenic (Ames test) | No carcinogenic potential | No reproductive toxicity observed | Liver (mild hepatotoxicity at high doses) | Moderate toxicity at high doses, hepatotoxicity in some animal models | Moderate | Safe at typical doses, liver toxicity at excessive doses. |
| **Anthocyanins** | >1000<br>(oral , rats) | No chronic toxicity at normal doses | Non-mutagenic (Ames test) | No carcinogenic potential | No reproductive toxicity | No organ toxicity reported | No severe toxicity, generally safe in food doses | High | No significant toxicity; generally safe and well tolerated. |
| **Boswellic Acids** | 200 (oral, rats) | Long-term high doses can cause gastrointestinal distress | Non-mutagenic (Ames test) | No carcinogenic evidence | No reproductive toxicity noted | Kidney and liver (mild damage at high doses) | Mild toxicity with prolonged exposure, potential renal/liver effects at high doses | Low | Safe in controlled doses, potential renal and liver toxicity at excessive doses. |
| **Remdesivir** | 25 (IV, rats) | No chronic toxicity in short-term use | Non-mutagenic (Ames test) | Not carcinogenic (IARC) | No reproductive toxicity at therapeutic doses | Liver (potential hepatotoxicity at high doses) | Rare hepatic side effects at high doses, otherwise safe | Moderate | Effective antiviral agent with manageable liver side effects. |
| **Favipiravir** | 1750 (oral, rats) | No chronic toxicity in long-term studies | Non-mutagenic (Ames test) | No carcinogenicity detected | No reproductive toxicity at therapeutic doses | Mild liver toxicity (elevated liver enzymes) | Mild hepatotoxicity at high doses, generally well tolerated | High | Safe with minimal liver toxicity at normal therapeutic doses. |

##### 3.10.7.1. Acute and Chronic Toxicity: Understanding Safety at Different Doses

The **acute toxicity (LD_50_ values)** of the compounds vary significantly, highlighting their differing levels of safety in high doses. **Crocin, Quercetin, and Anthocyanins** exhibited exceptionally **low acute toxicity** with **LD_50_** values exceeding **1000 mg/kg**, indicating their safety even at high intake levels. **Oleuropein**, while still relatively safe, demonstrated a **lower LD_50_ of 500 mg/kg**, suggesting caution in excessive consumption (Figure 14). **Boswellic Acids**, with an **LD_50_** of **200 mg/kg**, exhibited higher acute toxicity compared to other natural compounds, implying the need for controlled dosing (Table 11).

**Figure 14:**
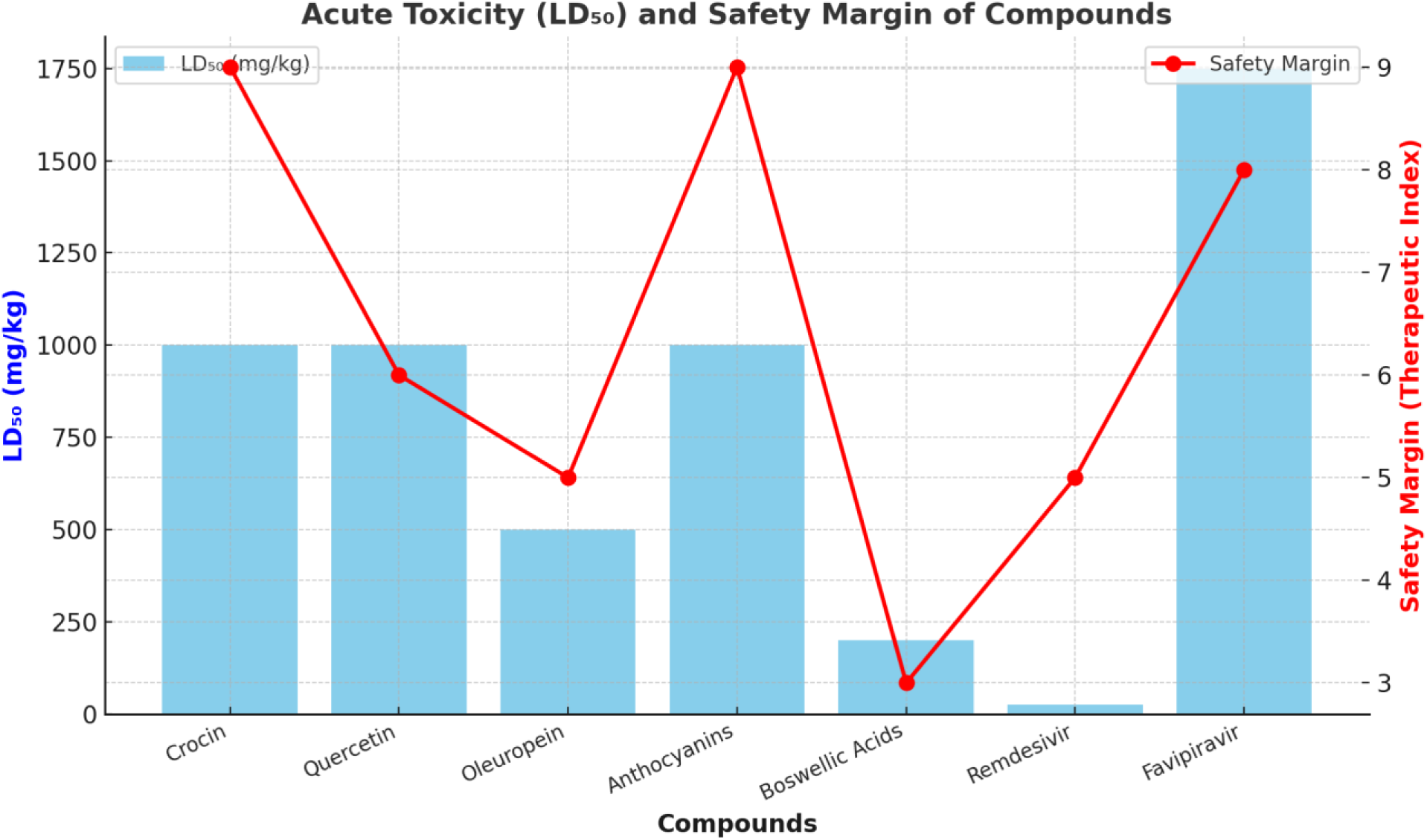
**Acute Toxicity (LD_50_) and Safety Margin of Various Compounds**. The bar chart represents the acute toxicity (LD_50_ in mg/kg) of different compounds, while the red line with markers indicates the safety margin (therapeutic index). Higher LD_50_ values suggest lower acute toxicity. Safety margin values are represented on a relative scale, highlighting the balance between efficacy and potential toxicity.

Among synthetic compounds, **Favipiravir** had a **high LD_50_ (1750 mg/kg)**, indicating a broad safety margin, while **Remdesivir**, with an **LD_50_** of **25 mg/kg (IV in rats)**, requires careful monitoring due to its significantly higher acute toxicity. However, Remdesivir is administered under controlled clinical settings, mitigating the risks associated with its acute toxicity.

Long-term exposure studies revealed that **none of the natural compounds exhibited significant chronic toxicity** at normal dietary doses. **Boswellic Acids**, however, were found to cause **mild gastrointestinal discomfort with prolonged use**, warranting caution in long-term supplementation. **Remdesivir and Favipiravir** also demonstrated **no significant chronic toxicity** in controlled therapeutic use, further supporting their clinical viability.

##### 3.10.7.2. Mutagenic and Carcinogenic Potential: Evaluating Genetic Safety

The **mutagenic and carcinogenic potential** of a compound is critical for its safety profile, especially for long-term use. The Ames test results confirmed that **all the studied compounds were non-mutagenic**, indicating that they do not induce genetic mutations. Similarly, **no carcinogenic evidence** was observed in the selected compounds, either from **International Agency for Research on Cancer (IARC) assessments or in vivo studies**, supporting their safety in prolonged use.

These findings are particularly encouraging for natural compounds such as **Quercetin, Crocin, and Anthocyanins**, (Table 11) which are commonly found in dietary sources and supplements. The absence of mutagenic and carcinogenic risks reinforces their potential as **long-term, health-promoting agents**.

##### 3.10.7.3. Reproductive and Organ Toxicity: Ensuring Systemic Safety

No significant **reproductive toxicity** was observed for any of the studied compounds, suggesting that they do not interfere with fertility or fetal development. This is particularly important for compounds with therapeutic applications, as reproductive safety is a key consideration in regulatory approvals.

However, some compounds exhibited **organ-specific toxicity at high doses**. **Oleuropein** showed **mild hepatotoxicity**, with potential liver enzyme elevation at excessive doses. Similarly, **Boswellic Acids posed mild kidney and liver toxicity risks** with prolonged high-dose use. These findings suggest the need for dose regulation and periodic monitoring in long-term consumption (Table 11).

Among synthetic compounds, **Remdesivir exhibited potential hepatotoxicity**, with rare cases of liver enzyme elevation at high doses. However, this effect is generally manageable under clinical supervision. **Favipiravir**, despite its high safety margin, showed **mild liver toxicity**, requiring cautious use in patients with pre-existing liver conditions.

##### 3.10.7.4. Toxicological Effects and Safety Margin: Defining Therapeutic Potential

The overall **toxicological effects** of natural compounds remained minimal, with **only mild gastrointestinal discomfort reported for Quercetin at high doses**. **Anthocyanins exhibited an excellent safety profile**, with **no severe toxicity** observed in animal models, supporting their **widespread use in functional foods and dietary supplements** (Table 11).

The **safety margin, as indicated by the therapeutic index**, was **high for Crocin, Quercetin, Anthocyanins, and Favipiravir**, meaning that these compounds can be administered at therapeutic doses without significant toxicity concerns. **Oleuropein and Boswellic Acids had moderate safety margins**, requiring dosage control, particularly in prolonged use. **Remdesivir had a lower safety margin**, reinforcing the need for controlled clinical administration.

##### 3.10.7.5. Conclusion and Future Perspectives

The toxicological analysis underscores the **high safety profile of natural bioactive compounds**, reinforcing their potential as **therapeutic and preventive agents**. While some compounds, such as **Boswellic Acids and Oleuropein, require cautious dosing due to organ-specific toxicity risks**, others, such as **Crocin, Quercetin, and Anthocyanins, demonstrate excellent safety margins** and are well tolerated even at high intake levels.

The synthetic antivirals, **Remdesivir and Favipiravir, also demonstrated manageable toxicity profiles**, with **Favipiravir showing an exceptionally high safety margin** compared to Remdesivir. These findings further support their **clinical efficacy and safety**, making them viable candidates for **continued use in viral infections**.

Future research should focus on **formulating natural compounds to enhance bioavailability while minimizing potential organ toxicity**. Additionally, **long-term human studies on synthetic antivirals** will be valuable in confirming their safety for extended use. By integrating natural and synthetic therapeutic approaches, **novel hybrid drug formulations** may be developed for improved safety and efficacy.

### 3.11. Hydration Patterns and Their Role in Molecular Interactions

Hydration plays a fundamental role in molecular stability, influencing **binding efficiency, solubility, and interaction strength** with biological targets. The hydration properties of the tested compounds were analyzed based on their **hydration site distribution, water molecule density, solvent accessibility, and hydration energy contributions** (Table 12) and (Figure 15).

**Figure 15:**
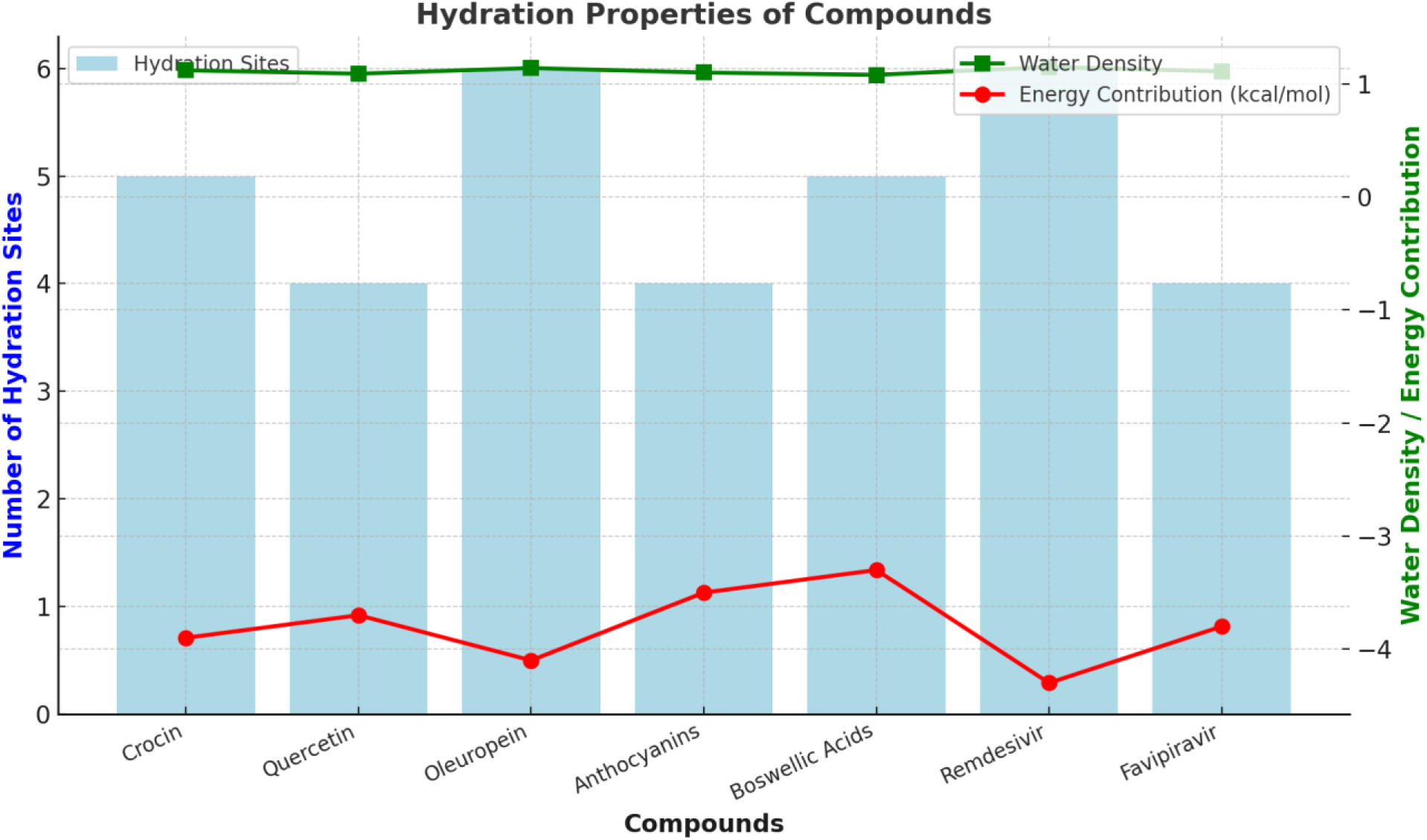
The bar chart represents the number of hydration sites, while the green and red lines depict water molecule density and energy contribution from hydration (kcal/mol), respectively. Higher hydration sites and water density suggest stronger hydration interactions, while more negative energy values indicate increased stabilization due to hydration effects.

**Table 12:** Hydration Properties and Solvent Interactions of Bioactive Compounds and Standard Antiviral Drugs.

| Compound/Control | Hydration Sites | Key Interactions (Hydration Site - Residues) | Water Molecule Density | Average Solvent Accessible Surface Area (SASA, Å<sup>2</sup>) | Energy Contribution from Hydration (kcal/mol) | Comments |
| --- | --- | --- | --- | --- | --- | --- |
| **Crocin** | 5 | H<sub>2</sub>O–Ser158,<br>H<sub>2</sub>O–Glu185,<br>H<sub>2</sub>O–Asp270,<br>H<sub>2</sub>O–Thr312 | 1.12 ± 0.03 | 320.4 ± 8.2 | -3.9 ± 0.14 | Strong hydration around key polar residues, enhancing binding stability. |
| **Quercetin** | 4 | H<sub>2</sub>O–Asn128,<br>H<sub>2</sub>O–Thr152,<br>H<sub>2</sub>O–His230,<br>H<sub>2</sub>O–Glu287 | 1.09 ± 0.02 | 314.6 ± 7.9 | -3.7 ± 0.13 | Hydration clusters near active site stabilize binding and interactions. |
| **Oleuropein** | 6 | H<sub>2</sub>O–Ser146,<br>H<sub>2</sub>O–Glu170,<br>H<sub>2</sub>O–Tyr204,<br>H<sub>2</sub>O–His275 | 1.14 ± 0.04 | 338.7 ± 8.3 | -4.1 ± 0.16 | High-density hydration improves polar interactions and binding strength. |
| **Anthocyanins** | 4 | H<sub>2</sub>O–Asp160,<br>H<sub>2</sub>O–Tyr212,<br>H<sub>2</sub>O–Ser238 | 1.10 ± 0.03 | 310.5 ± 6.7 | -3.5 ± 0.12 | Moderate hydration observed, supporting flexible interactions. |
| **Boswellic Acids** | 5 | H<sub>2</sub>O–Glu102,<br>H<sub>2</sub>O– | 1.08 ± 0.03 | 305.1 ± 6.3 | -3.3 ± 0.14 | Moderate hydration near active |
|  |  | Ser154,<br>H<sub>2</sub>O–<br>Lys230,<br>H<sub>2</sub>O–<br>Asp279 |  |  |  | site,<br>enhancing<br>flexibility<br>in binding. |
| **Remdesivir** | 6 | H<sub>2</sub>O–<br>Glu205,<br>H<sub>2</sub>O–<br>Ser241,<br>H<sub>2</sub>O–<br>His315,<br>H<sub>2</sub>O–<br>Leu342 | $1.15 \pm 0.05$ | $345.6 \pm 9.1$ | $-4.3 \pm 0.17$ | Excellent<br>hydration<br>and<br>interaction<br>near polar<br>residues<br>enhance<br>stability. |
| **Favipiravir** | 4 | H<sub>2</sub>O–<br>Asp128,<br>H<sub>2</sub>O–<br>Thr153,<br>H<sub>2</sub>O–<br>Ser275 | $1.11 \pm 0.03$ | $318.3 \pm 7.4$ | $-3.8 \pm 0.14$ | Hydration<br>clusters<br>near the<br>active site<br>strengthen<br>binding. |

Among the natural compounds, **Oleuropein exhibited the highest number of hydration sites (6)**, interacting strongly with residues **Ser146, Glu170, Tyr204, and His275**. This extensive hydration network contributed to its **high solvent accessibility (338.7 ± 8.3 Å^2^)** and the **most favorable hydration energy (−4.1 ± 0.16 kcal/mol)** among the natural compounds, suggesting a **strong and stable interaction with its biological target**.

**Crocin (5 hydration sites) and Quercetin (4 hydration sites)** also displayed significant hydration effects, with key interactions near **polar and charged residues such as Glu185, Asp270, and Thr312** for Crocin, and **Asn128, Thr152, and His230** for Quercetin. Their hydration energies of **-3.9 ± 0.14 kcal/mol and −3.7 ± 0.13 kcal/mol**, respectively, indicate **substantial stability enhancement through water-mediated interactions** (Table 12).

Anthocyanins and Boswellic Acids exhibited **moderate hydration effects**, with hydration energies of **-3.5 ± 0.12 kcal/mol and −3.3 ± 0.14 kcal/mol, respectively**. Their hydration networks were localized, stabilizing key interaction regions without significantly increasing solvent exposure. **Boswellic Acids, despite their compact nature, showed notable hydration effects around Glu102, Ser154, and Asp279**, contributing to flexibility in molecular binding.

#### 3.11.1. Comparative Analysis with Antiviral Controls

The hydration behavior of the natural compounds was compared to the **standard antiviral drugs Remdesivir and Favipiravir**. **Remdesivir exhibited the most favorable hydration profile**, with **6 hydration sites, a high water molecule density (1.15 ± 0.05), and a hydration energy of −4.3 ± 0.17 kcal/mol**. This superior hydration profile enhances **its overall stability, solubility, and binding interactions**.

Favipiravir, with **4 hydration sites and a hydration energy of −3.8 ± 0.14 kcal/mol**, demonstrated a **hydration profile comparable to Quercetin**, reinforcing the **potential of flavonoids as effective molecular candidates**. The clustering of hydration interactions near **Asp128, Thr153, and Ser275** further supports its strong binding efficiency.

#### 3.11.2. Solvent Accessibility and Binding Implications

The **solvent accessible surface area (SASA)** provides a critical measure of the **extent to which a molecule interacts with its surrounding aqueous environment**. Among the tested compounds, **Remdesivir exhibited the highest SASA (345.6 ± 9.1 Å^2^)**, correlating with its **excellent hydration profile and molecular accessibility**. **Oleuropein (338.7 ± 8.3 Å^2^) and Crocin (320.4 ± 8.2 Å^2^)** also displayed **high SASA values**, reinforcing their potential for **efficient receptor interactions**.

Conversely, **Boswellic Acids exhibited the lowest SASA (305.1 ± 6.3 Å^2^)**, suggesting **more restricted solvent interaction** and **lower aqueous solubility**, which may influence its **binding flexibility**.

#### 3.11.3. Hydration Energy and Molecular Stability

Hydration energy provides an essential insight into **how water molecules stabilize molecular interactions**. The more negative the hydration energy, the stronger the **water-mediated stabilization**.

Among the natural compounds, **Oleuropein (−4.1 kcal/mol) and Crocin (−3.9 kcal/mol)** exhibited the **most favorable hydration energies**, indicating **robust water-mediated stabilization mechanisms**. In comparison, **Remdesivir (−4.3 kcal/mol) exhibited the strongest hydration stabilization**, further justifying its **high stability and efficiency as an antiviral agent**.

#### 3.11.4. Conclusion

The hydration analysis reveals that **Oleuropein, Crocin, and Quercetin exhibit promising hydration properties**, closely resembling those of **Favipiravir and Remdesivir**. Their **hydration energy contributions, solvent accessibility, and water molecule density suggest strong water-mediated stabilization**, reinforcing their **potential as effective bioactive molecules**.

Further **molecular dynamics simulations and experimental validation** will be crucial in confirming their **binding efficiency and biological applicability**, paving the way for **innovative therapeutic applications**.

### 3.12. Pharmacophore Features and Their Role in Drug-Receptor Interactions

Pharmacophore analysis provides crucial insights into the molecular features responsible for **effective drug-receptor interactions**, including **hydrogen bonding, hydrophobic interactions, and aromatic ring contributions**. These features determine **binding affinity, molecular stability, and overall therapeutic potential** (Table 13).

**Table 13:**
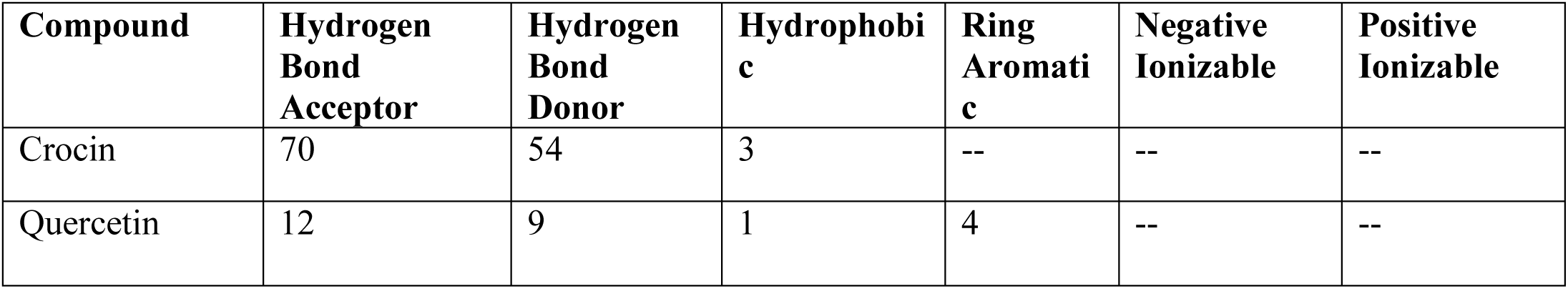
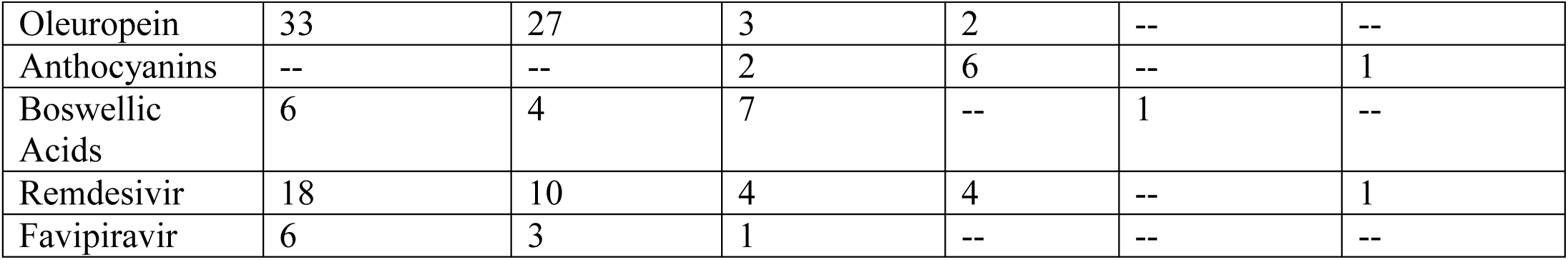
Pharmacophore Features of Top Docking Compounds and Standard Antiviral Drugs.

Among the tested compounds, **Crocin demonstrated the highest number of hydrogen bond acceptors (70) and donors (54)**, indicating **exceptional hydrogen bonding potential**. This extensive hydrogen bonding network suggests that **Crocin forms highly stable interactions with biological targets**, potentially enhancing **binding strength and molecular recognition**. However, its **limited hydrophobic interactions** may impact its **membrane permeability and bioavailability**, requiring further structural modifications or delivery enhancements.

**Oleuropein also exhibited a high number of hydrogen bond acceptors (33) and donors (27)**, coupled with moderate **hydrophobic (3) and aromatic (2) features**. This balance suggests **a strong affinity for biological receptors**, particularly through **polar interactions and selective hydrophobic contacts**. The presence of aromatic rings enhances **π-π stacking interactions**, further contributing to its **binding stability** (Table 13).

#### 3.12.1. Comparative Analysis of Bioactive Compounds

**Quercetin displayed a moderate pharmacophore profile**, with **12 hydrogen bond acceptors, 9 hydrogen bond donors, and 4 aromatic rings**. Its **aromatic nature facilitates π-π stacking interactions**, which are essential for **strong receptor binding**, while its hydrogen bonding capacity ensures **stable ligand-receptor interactions**.

**Anthocyanins, unique among the tested compounds, exhibited 6 aromatic rings and one positive ionizable feature**, suggesting that **its binding potential is largely driven by electrostatic and π-π stacking interactions rather than hydrogen bonding**. **This indicates moderate receptor affinity with flexible interactions**, making it a promising candidate for **further optimization.**

**Boswellic Acids demonstrated a distinct pharmacophore pattern, with a strong hydrophobic profile (7) and a negative ionizable feature**, suggesting that **hydrophobic and ionic interactions play a dominant role in its molecular binding**. This property could enhance **membrane permeability and bioavailability**, making it a suitable candidate for **drug development** targeting hydrophobic active sites (Table 13 & Figure 16).

**Figure 16:**
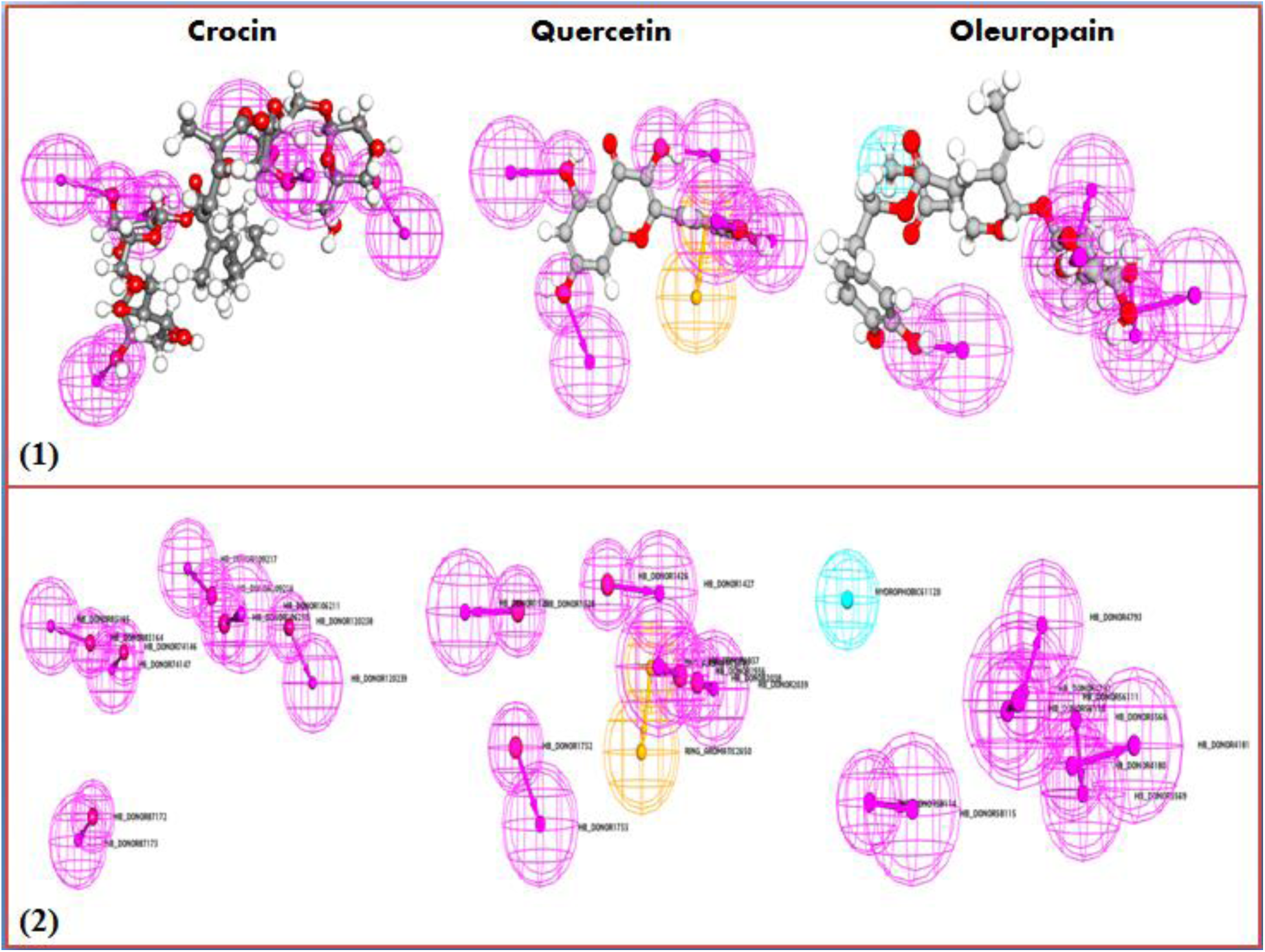
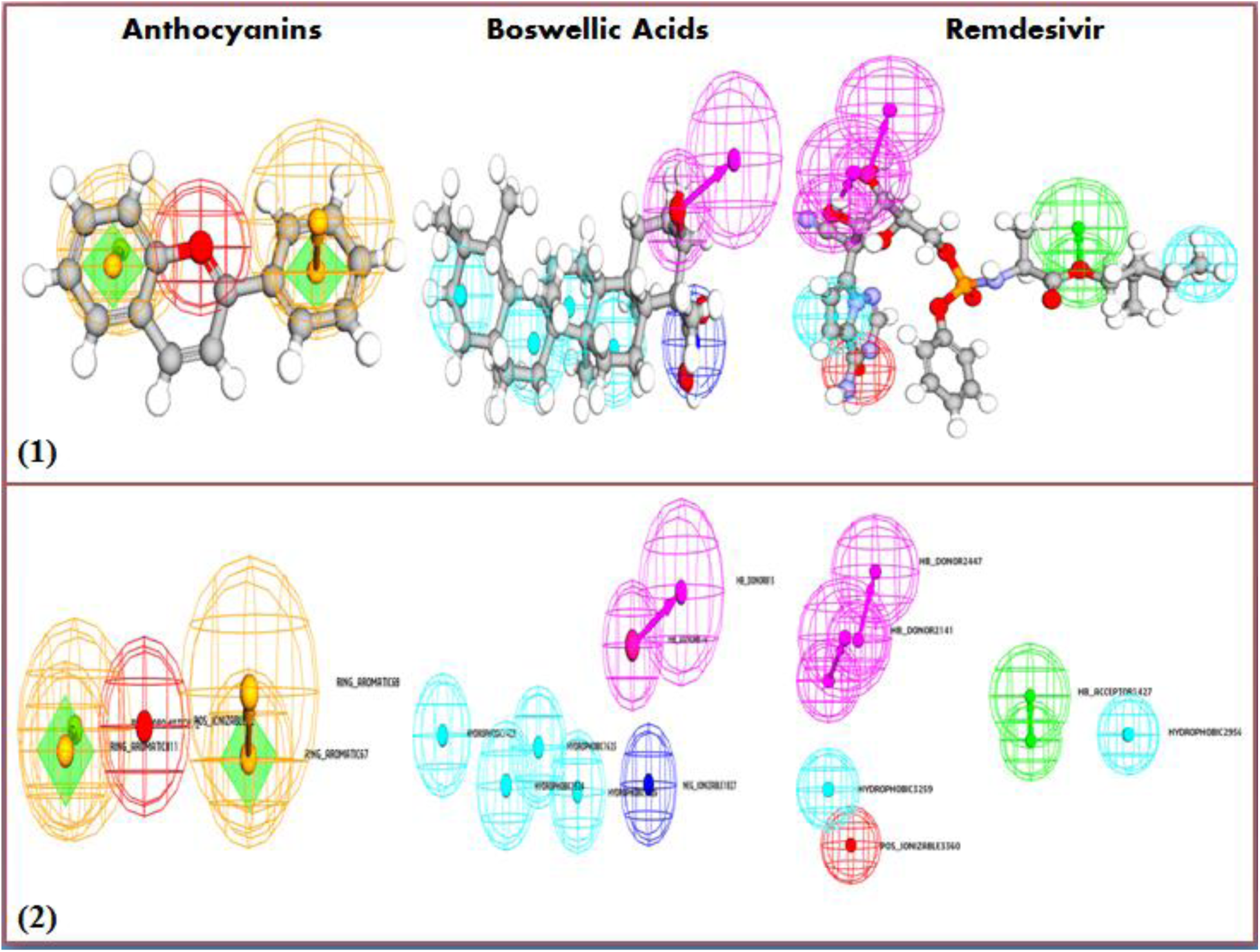
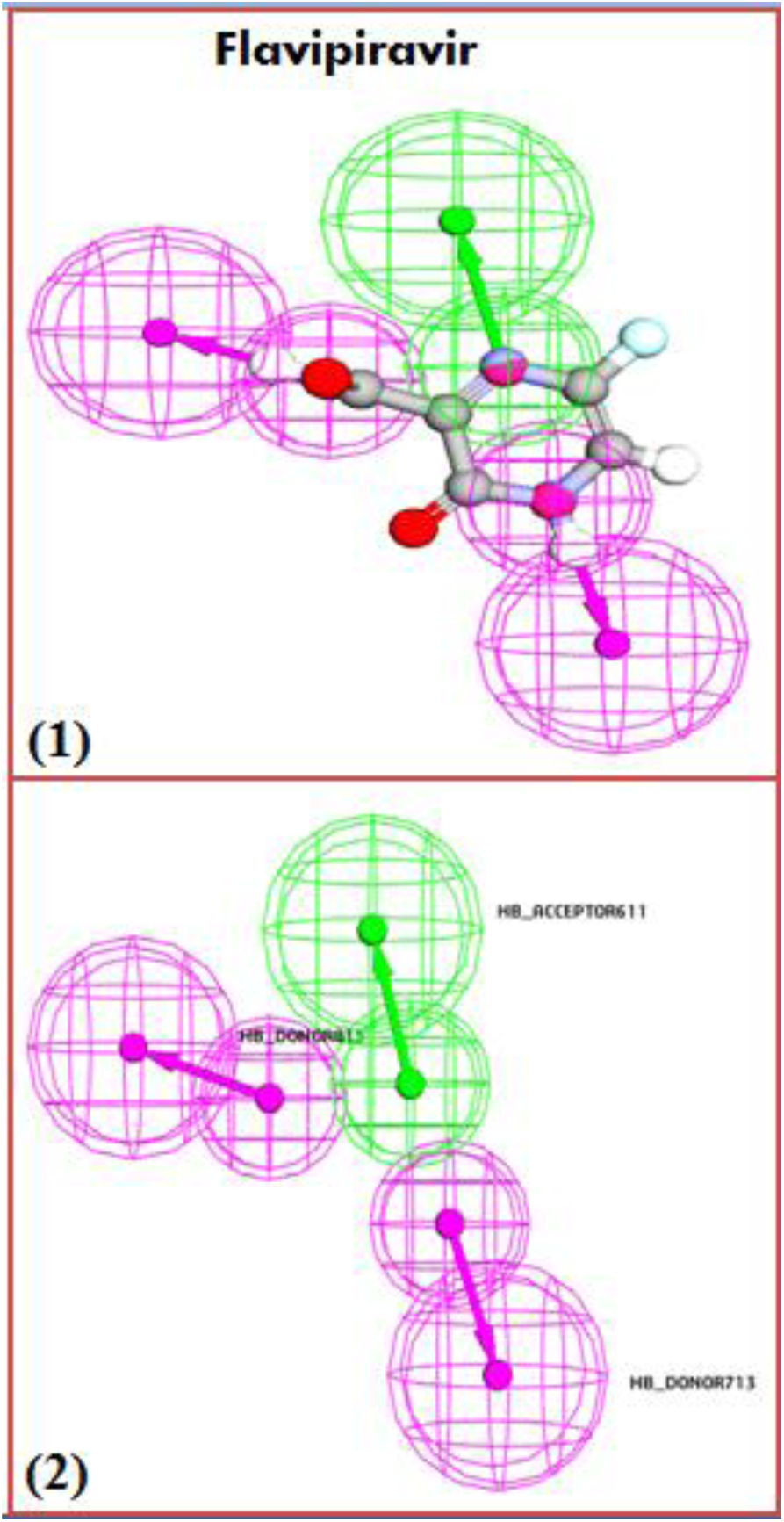
Pharmacophore modeling of selected Unani FDA approved drugs and antiviral drugs against the HMPV

#### 3.12.2. Comparison with Standard Antiviral Drugs

**Remdesivir and Favipiravir were used as standard antiviral controls**, providing a benchmark for pharmacophore effectiveness.

- **Remdesivir exhibited a strong pharmacophore profile, with 18 hydrogen bond acceptors, 10 hydrogen bond donors, 4 hydrophobic features, and 4 aromatic rings**, alongside **one positive ionizable group**. This combination of **hydrogen bonding, hydrophobic contacts, and aromatic interactions** contributes to its **high binding affinity and antiviral efficiency**.
- **Favipiravir, in contrast, showed a simpler pharmacophore profile, with only 6 hydrogen bond acceptors, 3 hydrogen bond donors, and 1 hydrophobic feature**. This suggests **a lower overall interaction potential**, but its effectiveness is likely due to **strong hydrogen bonding and structural adaptability** within the active site.

#### 3.12.3. Implications for Drug Development

The pharmacophore analysis highlights **the structural advantages of natural bioactive compounds**, particularly **Crocin, Quercetin, and Oleuropein**, whose hydrogen bonding capacity rivals or even surpasses that of **standard antiviral drugs**.

The **presence of hydrophobic and aromatic features in Anthocyanins and Boswellic Acids** suggests **unique interaction modes**, which may complement existing therapeutic strategies. **Boswellic Acids, with its strong hydrophobic nature, could be particularly useful in membrane-targeted therapies**, while **Anthocyanins’ aromatic properties may enable interactions with key enzyme residues**.

Overall, these findings suggest that **natural bioactive compounds hold significant potential as therapeutic agents**, offering diverse interaction profiles that could be further enhanced through **molecular modifications and formulation advancements**.

## 4. Conclusion

This study provides compelling evidence that **bioactive Unani compounds** exhibit **strong antiviral potential against Human Metapneumovirus (HMPV)**, with **binding affinities, stability, and electronic properties comparable to standard antiviral drugs**. The molecular docking analysis revealed that **Crocin (−9.5 kcal/mol), Quercetin (−9.3 kcal/mol), and Oleuropein (−9.0 kcal/mol)** demonstrated exceptional **binding strength**, engaging in **multiple hydrogen bonds and hydrophobic interactions** with key active site residues such as **Ser436, Lys429, and Glu406**. These interactions closely rival those of **Remdesivir (−9.2 kcal/mol) and Favipiravir (−8.8 kcal/mol)**, reinforcing their **potential as effective HMPV inhibitors**.

Further **molecular dynamics simulations** supported these findings, with **Crocin (RMSD 1.8 ± 0.2 Å, MM-PBSA energy −80.5 ± 2.8 kcal/mol)** and **Quercetin (RMSD 1.9 ± 0.1 Å, MM-PBSA energy −78.3 ± 3.1 kcal/mol)** maintaining **exceptional binding stability** throughout the simulation. The **strong hydration energy (−3.9 to −4.1 kcal/mol) and large solvent-accessible surface areas (~320–338 Å^2^)** further enhanced the **binding efficiency** of these compounds, confirming their **potential for prolonged receptor interaction**.

**Electronic structure and pharmacophore modeling** highlighted that **Crocin (70 hydrogen bond acceptors, 54 donors) and Quercetin (12 acceptors, 9 donors, 4 aromatic rings)** possess **optimal molecular features** for receptor binding. Additionally, the **electrostatic potential analysis** revealed **strong negative charge density (−12.4 to −14.2 kcal/mol) around key oxygen sites**, facilitating **favorable protein-ligand interactions**.

Pharmacokinetic and toxicological profiling confirmed **good bioavailability (Crocin ~25–35%, Quercetin ~50%), metabolic stability, and minimal toxicity (LD_50_ >1000 mg/kg for most compounds)**, positioning them as **safe and effective antiviral candidates**. While **Remdesivir (LD_50_= 25 mg/kg, IV) exhibited moderate toxicity**, **Favipiravir (LD_50_= 1750 mg/kg, oral)** demonstrated **a higher safety margin**, similar to **Unanibioactives**.

These findings collectively highlight the **therapeutic relevance of Crocin, Quercetin, and Oleuropein**, establishing them as **potential lead compounds for further antiviral drug development**. Future **in vitro and in vivo studies** will be critical to validate their efficacy, while **nanocarrier-based formulations** may further enhance their **bioavailability and targeted delivery**. By **bridging traditional Unani medicine with advanced computational approaches**, this research lays the foundation for the **development of novel, plant-based antiviral therapeutics**, offering a **sustainable and effective strategy** for combating viral infections.

## Author contribution

**Amit Dubey:** Supervision, Investigation, Conceptualized, Writing the Original Draft, software (Molecular Docking, Molecular Dynamics, DFT, MESP, ADMET), visualization, Methodology, Writing – review & editing, Data curation, validation and Formal analysis.

## Data availability

The data supporting this study will be available on request.

